# PI3Kγ inhibition suppresses microglia/TAM accumulation in glioblastoma microenvironment to promote exceptional temozolomide response

**DOI:** 10.1101/2020.05.14.097121

**Authors:** Jie Li, Megan M. Kaneda, Jun Ma, Ming Li, Kunal Patel, Tomoyuki Koga, Aaron Sarver, Frank Furnari, Beibei Xu, Sanjay Dhawan, Jianfang Ning, Hua Zhu, Anhua Wu, Gan You, Tao Jiang, Andrew S. Venteicher, Jeremy N. Rich, Christopher K. Glass, Judith A. Varner, Clark C. Chen

**Affiliations:** Department of Neurosurgery, University of Minnesota, Minneapolis, MN, USA; Department of Pharmacology and Pathology, University of California, San Diego, La Jolla, CA, USA; Department of Neurosurgery, University of California Los Angeles, Los Angeles, CA, USA; Institute for Health Informatics, University of Minnesota, Minneapolis, MN, USA; Ludwig Institute for Cancer Research, San Diego, La Jolla, CA, USA; Department of Pediatrics, The First Hospital of China Medical University, Shenyang, China; Department of Neurosurgery, The First Hospital of China Medical University, Shenyang, China; Department of Neurosurgery, Beijing Tiantan Hospital, Capital Medical University, Beijing, China; Department of Molecular Neuropathology, Beijing Neurosurgical Institute, Capital Medical University, Beijing, China; Department of Medicine, Division of Regenerative Medicine, University of California, San Diego,, La Jolla, CA, USA; Department of Medicine, University of California, San Diego, La Jolla, CA, USA

**Keywords:** glioblastoma, exceptional responders, IL11, PI3Kγ, microglia/macrophages

## Abstract

Precision medicine in oncology leverages clinical observations of exceptional response. Towards an understanding of the molecular features that define this response, we applied an integrated, multi-platform analysis of RNA profiles derived from clinically annotated glioblastoma samples. This analysis suggested that specimens from exceptional responders are characterized by decreased accumulation of microglia/macrophages in the glioblastoma microenvironment. Glioblastoma-associated microglia/macrophages secreted interleukin 11 (IL11) to activate STAT3-MYC signaling in glioblastoma cells. This signaling induced stem cell states that confer enhanced tumorigenicity and resistance to the standard-of-care chemotherapy, temozolomide (TMZ). Targeting a myeloid cell restricted isoform of PI3K, PI3Kγ, by pharmacologic inhibition or genetic inactivation, disrupted this signaling axis by suppressing microglia/macrophage accumulation and associated IL11 secretion in the tumor microenvironment. Mirroring the clinical outcomes of exceptional responders, PI3Kγ inhibition synergistically enhanced the anti-neoplastic effects of TMZ in orthotopic murine glioblastoma models. Moreover, inhibition or genetic inactivation of PI3Kγ in murine glioblastoma models recapitulated expression profiles observed in clinical specimens isolated from exceptional responders. Our results suggest key contributions from tumor-associated microglia/macrophages in exceptional responses and highlight the translational potential for PI3Kγ inhibition as a glioblastoma therapy.

**Significance Statement:** Understanding the basis for exceptional responders represents a key pillar in the framework of precision medicine. In this study, we utilized distinct informatics platforms to analyze the expression profiles of clinically annotated tumor specimens derived from patients afflicted with glioblastoma, the most common form of primary brain cancer. These analyses converged on prognostic contributions from glioblastoma-associated microglia/macrophages. Glioblastoma-associated microglia secreted interleukin 11 (IL11) to activate a STAT3-MYC signaling axis in glioblastoma cells, facilitating resistance to the standard-of-care chemotherapy, temozolomide. Microglia recruitment and IL11 secretion were dependent on the myeloid specific phosphoinositide-3-kinase gamma isoform (PI3Kγ). Inhibition or genetic inactivation of PI3Kγ in murine glioblastoma models recapitulated expression profiles observed in specimens derived from exceptional responders, suggesting potential for clinical translation.

## Introduction

Key frameworks in cancer biology have been built on insights gained through genetic analysis of rare, hypo-morphic mutations/epigenetic events that render hypersensitivity to specific experimental conditions (1). The study of exceptional responders can be viewed in this light, as the tumors in these patients are often characterized by genetic or epigenetic events that confer exquisite sensitivity to the treatment rendered. Most studies define exceptional responders as the <10% of patients who markedly respond to a therapy that confer little benefit to the remaining population (2). Here, we adopt this definition and study the <10% of patients with glioblastoma (World Health Organization grade IV glioma), the most common form of primary brain cancer in adults, who survive beyond two years after the standard-of-care treatment with concurrent radiation and temozolomide (TMZ).

Previous studies of exceptional glioblastoma responders have yielded two key observations. First, while glioblastomas harboring isocitrate dehydrogenase (IDH) mutations are generally associated with improved survival (3), such mutations are not required for long-term survivorship (4, 5). Second, in independent studies, clinical specimens from long-term survivors are more likely to harbor promoter methylation attenuating the expression of O6-methylguanine-DNA-methyltransferase (MGMT), a DNA repair protein that confers TMZ resistance (6–9). These results suggest the critical contribution of TMZ efficacy to long-term survivorship. Notably, FDA approval of TMZ for glioblastoma factored into consideration the small “tail” of patients who survived beyond two years, since the treatment-associated gain in median survival was minimal (10, 11). Beyond these observations, there is little commonality in the various studies of long-term glioblastoma survivorship (12). A central limitation in this literature is that key clinical factors that dominate clinical survival prognostication for glioblastoma patients, such as patient age, neurologic function, and extent of surgical resection (13, 14), are rarely considered in efforts to develop survival gene signatures.

Studies of long-term glioblastoma survivorship have largely focused on interrogating biological processes intrinsic to the tumor cell (12). However, 30-50% of the cells in clinical glioblastoma specimens are non-neoplastic, consisting predominantly of microglia or Gr1+ macrophages (15). Importantly, glioblastoma cells are indelibly influenced by these non-neoplastic cells. Microglia are progenies of primitive yolk sac myeloid precursors that reside in the central nervous system (CNS) (16). Macrophages are derived from bone marrow myeloid precursors that circulate in the peripheral blood and enter the brain during tumor development; some differentiate into immune suppressive, Gr1+ tumor-associated macrophages (TAM) (17). While microglia and TAM can be studied as distinct entities in murine glioblastoma models (18), dissociating the contributions of these distinct myeloid progenies is difficult in studies of human clinical glioblastoma specimens (19). As such, microglia/TAM are often grouped into a single entity in human studies.

Many non-CNS tumors secrete chemokines to attract peripheral myeloid cells. These chemokines trigger activation of phosphoinositide-3-kinase gamma isoform (PI3Kγ) in myeloid cells to trigger chemotaxis (20) and recruitment into the tumor microenvironment (21, 22). In mammalian cells, there are three classes of PI3K: Class I PI3K generates secondary messengers to regulate cell growth, movement, and differentiation (23). Classes II and III PI3K play major roles in endocytosis/endosome trafficking (24–26). There are four class I PI3K isoforms (PI3Kα, β, and δ). The current models suggest that PI3Kα, β, and δ(classified as class IA) are primarily activated downstream of receptor tyrosine kinases while PI3Kγ (classified as class IB) is primarily activated in response to G-protein coupled receptors (GPCR) (27, 28), but can also be activated by receptor tyrosine kinases such as VEGFR (21). While PI3Kγ activation is critical for recruitment of TAM to non-CNS tumors (21, 22, 29–33), the relevance of this finding to glioblastoma recruitment of microglia or TAM remains unclear.

Dynamic and complex interactions occur between glioblastoma and microglia/TAM in the tumor microenvironment. Non-transformed microglia isolated from surgical specimens derived from epilepsy patients as well as macrophages cultured from peripheral human blood suppressed glioblastoma tumorigenicity (34), suggesting that native microglia/macrophage serve key roles in immune surveillance against cancer. This surveillance function is lost in microglia/TAM isolated from the murine and human glioblastomas (35). Instead, these microglia/TAM secrete a multitude of growth factors (36) and cytokines, including interleukin-6 (IL6) to promote glioblastoma growth (37). RNAseq profiling of glioblastoma-associated microglia/TAM revealed distinct sub-populations exhibiting gene expression patterns distinct from native microglia/TAM (38). Notably, single-gene biomarkers intended to proxy cell states in microglia/macrophage are often altered for reasons unrelated to the proxied cell state (39, 40). Reflecting this complexity, there is a notable incongruence in studies correlating clinical survival to the expression of single genes thought to reflect microglia/TAM abundance or cell-state, such as the expression of CD204 or ionized calcium-binding adaptor molecule 1 (Iba1) (41). These observations underscore the need for an integrated informatics approach (42, 43).

Here, we used independent informatics approaches to understand the molecular basis underlying exceptional response after standard-of-care glioblastoma treatment and uncovered contribution from glioblastoma-associated microglia/TAM. Glioblastoma-associated microglia secrete high levels of the pro-inflammatory cytokine IL11, which in turn, triggers the STAT3-MYC signaling axis in glioblastoma cells to induce stem-cell states that confer therapeutic resistance. Pharmacologic inhibition or genetic inactivation of PI3Kγ suppresses microglia/TAM accumulation in the tumor microenvironment to recapitulate expression signatures observed in clinical specimens derived from exceptional responders. These findings suggest PI3Kγ inhibition as a promising strategy for glioblastoma therapy.

## Results

### Gene signatures implicate contribution of glioblastoma-associated microglia/macrophages to clinical response

Realizing the hazard of “chance” association between random gene signatures and clinical survival (44), we began our analysis by interrogating published glioblastoma survival signatures to determine whether the association of gene expression signatures and survival can be recapitulated in independent patient cohorts. For this analysis, we focused on glioblastoma with wild-type IDH (wtIDH), given the paucity of information on survival prognostication in this patient population (45). From the literature, we identified 18 published glioblastoma survival gene signatures (Table S1) and tested them using the clinically annotated expression profiles derived from wtIDH patients in The Cancer Genome Atlas (TCGA, http://cancergenome.nih.gov/) and the REMBRANDT glioblastoma databases (46). Of the 18 published signatures, only two, Kim et al (47) and Lewis et al (48), consistently showed survival association in both patient cohorts (Table S1). Lowered expression levels of these gene signatures were associated with improved survival.

Clinical variables such as age, Karnofsky Performance Score (KPS, a clinical scale commonly used to assess a patient’s functional status), extent of resection, as well as temozolomide (TMZ) treatment potently influence patient survival (49). To assess whether survival association of the two gene signatures persist after controlling for these clinical variables, we combined the two signatures into an 82 gene panel (Table S2) and tested for survival association using a matched cohort design. We identified a total of ten patients in the Chinese Glioma Genome Atlas (CGGA) database who survived >2 years after gross total resection and TMZ treatment (~10% of the cohort). Using single sample gene set enrichment analysis (ssGSEA), we then compared the cumulative expression of these 82 genes in the >2 year survivors relative to a cohort of ten age- and KPS-matched wild-type glioblastoma patients who survived <1 year after gross total resection and TMZ treatment. The expression score of the combined 82 gene signature was lower in the >2 year survivors relative to the <1 year survivors (*p*=0.036, Fig. 1A). Functional annotation (50) of the combined signature revealed enrichment for genes involved in wound healing, inflammatory processes, and negative regulation of proliferation (Fig. 1B, and Table S2).

**Fig. 1.**
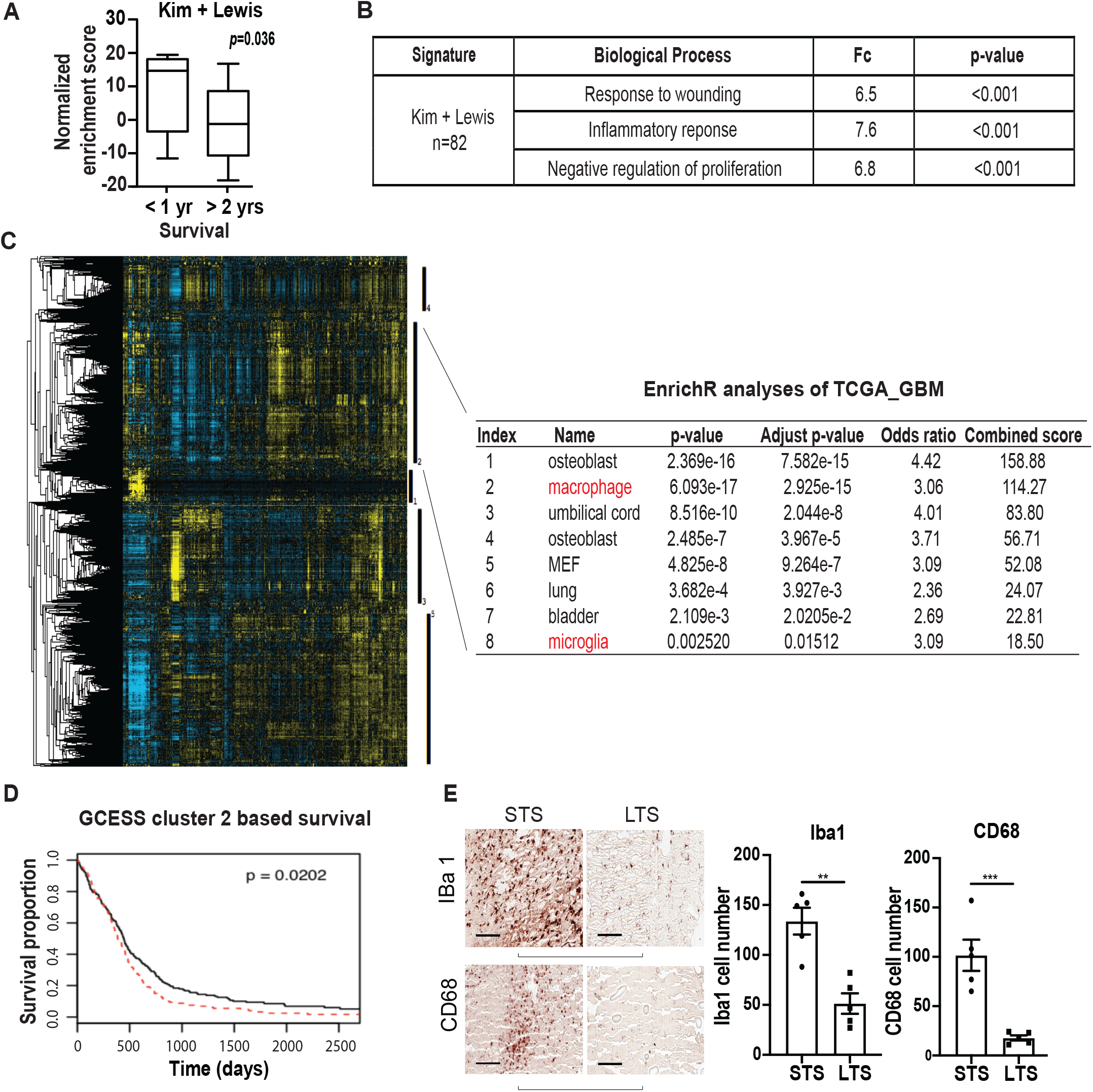
Survival analysis implicates contribution of glioblastoma-associated macrophages/microglia to clinical survival. (A) ssGSEA analysis demonstrating that the ssGSEA expression score of the 82 survival-associated genes (combining the Kim and Lewis signatures) was significantly lower in the >2 year survivors relative to the <1 year survivors. These cohorts were matched for age and KPS. All patients underwent gross total resection and TMZ treatment. (B) Shown are the pathways enriched for the 82 survival associated genes. (C, D) Gene Cluster Expression Summary Score (GCESS) analysis of TCGA glioblastomas. Cluster of genes that are 1) highly correlated to one another and 2) associated with overall survival are shown as heat map; each column represents a gene and each row a patient. The five survival associated gene clusters are indicated by the vertical black bars (to the right of the heat map). Macrophage/microglia gene signatures are enriched in cluster 2 (C). Higher expression of genes in cluster 2 was associated with poor clinical survival (D). High and low expression was defined based on median expression value. n=539; p=0.0202. (E) Immunohistochemical staining of macrophage and microglia markers, Iba1 and CD68, in CGGA patient cohort of long-term survival (LST) and matched short-term survivals (STS). Each row represents an LST with a matched STS. Two representative matched pairs are shown. The statistics were derived from five matched pairs. **, p<0.01; ***, p<0.001.

In parallel, we performed a survival analysis of the TCGA and REMBRANDT datasets using Gene Cluster Expression Summary Score (GCESS) (51), to identify gene clusters whose expression are 1) highly correlated to one another and 2) associated with overall survival in glioblastoma patients. This unbiased analysis revealed gene clusters implicating the contribution of microglia and TAM to clinical survival in both the TCGA (Fig. 1C, 1D) and the REMBRANDT datasets (Fig. S1). Higher expression levels of these gene clusters were associated with poor survival.

Integrating insights from these two informatics analyses, we hypothesized that accumulation of glioblastoma-associated microglia/macrophage induced inflammatory responses that contributed to poor patient survival. To test this hypothesis, we stained histology specimens prepared from the matched CGGA patients for two microglia/macrophage markers, Iba1 and CD68. Supporting our hypothesis, lower levels of Iba1 and CD68 staining were consistently observed in the exceptional responders relative to the matched control (Fig. 1E).

### Microglia-secreted IL11 enhances glioblastoma tumorigenicity and TMZ resistance

The functional annotation shown in Fig. 1B implicates cell states resembling those of stem cells since they 1) play critical roles in wound healing (52), 2) generally exhibit slower growth kinetics (53), and 3) participate in the inflammatory response (54). Since glioblastoma cells with stem cell properties exhibit features associated with poor clinical outcome, including increased tumorigenicity (55) and therapeutic resistance (56), our analyses suggest that microglia/macrophage in the tumor microenvironment may induce glioblastoma stem cell states that contribute to shortened survival. Supporting this hypothesis, co-culturing of glioblastoma cells with a transformed human microglia cell line (hMG) enhanced the *in vitro* and *in vivo* tumorigenicity (Fig. S2A and S2B) of glioblastoma cells as well as their TMZ resistance (Fig. S2C).

Realizing that the physiology of *in vitro* cultured microglia may differ from their *in vivo* roles (57), we tested whether our observations could be recapitulated using murine microglia freshly isolated from a syngeneic glioblastoma model. To this end, murine glioblastoma-associated microglia (mMG_gl_, GFP-CD45^low^CD11b^+^Gr1^−^ cells, see gating strategy, Fig. S3) were isolated from murine glioblastomas formed after orthotopic implant of GFP-labelled GL261 cells. Supporting results derived from cultured hMG, co-culture of GL261 with freshly isolated mMGgl enhanced *in vitro* colony formation potency by ~ 3-fold (Fig. 2A). Further, intracranial co-implantation of GL261 with freshly isolated mMG_gl_ enhanced *in vivo* tumor growth (Fig. 2B) as demonstrated by the shortened survival. Tumor-promoting effects were not observed when GL261 cells were co-cultured or co-implanted with GFP^−^CD45^low^CD11b^+^Gr1^−^ microglia isolated from normal murine brains (mMG_nb_, Fig. 2A and 2B). Moreover, co-culturing with mMG_gl_ but not mMG_nb_, enhanced GL261 resistance to TMZ (Fig. 2C). Of note, similar effects were observed when glioblastoma-associated macrophages (defined by GFP^−^CD45^high^CD11b^+^Gr1^+^) were tested *in vivo* (Fig. S2D).

**Fig. 2.**
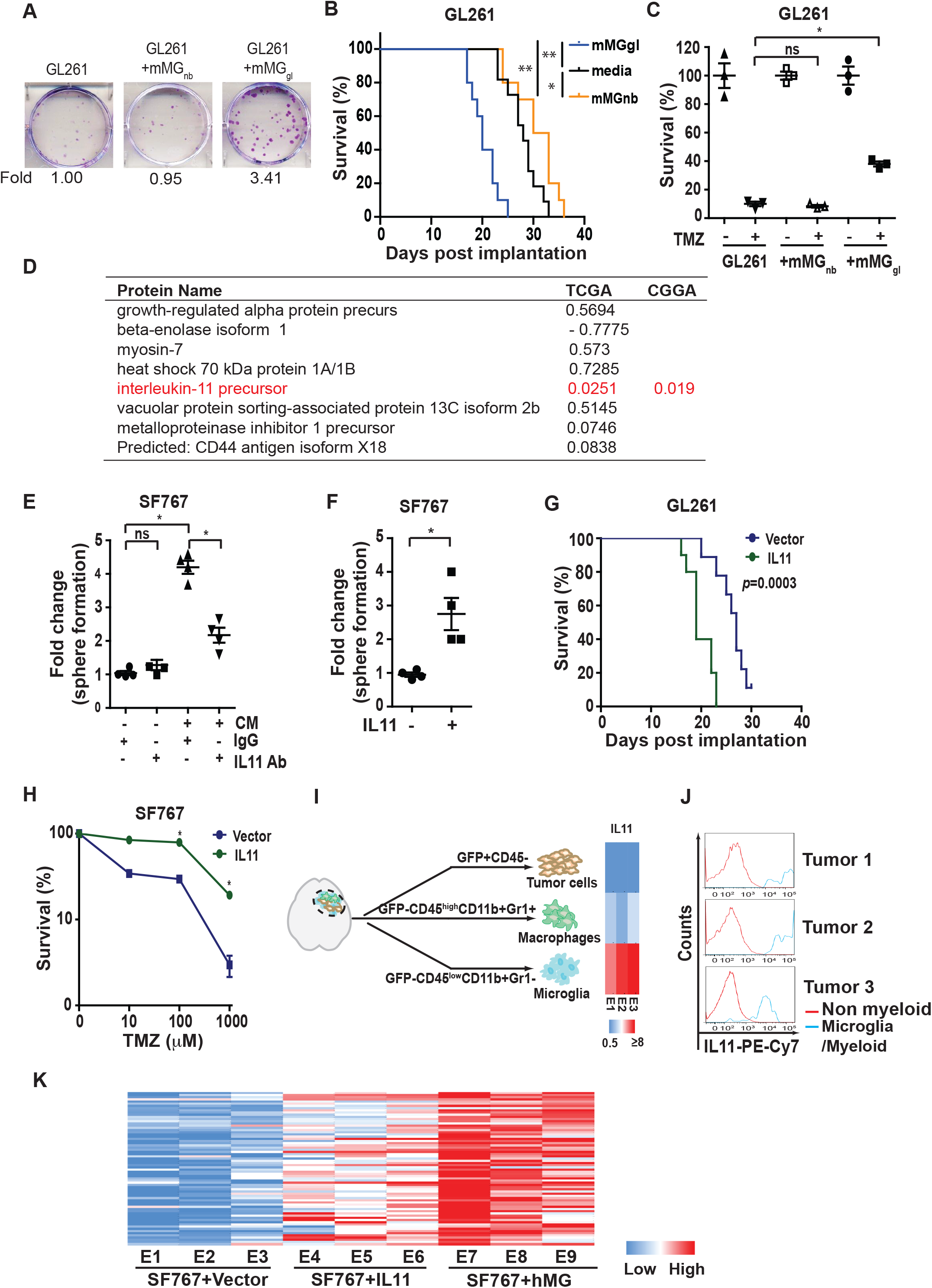
Glioblastoma-associated microglia release IL11 to enhance glioblastoma tumorigenicity and TMZ resistance. (A) Upper: Representative images of *in vitro* colony formation after co-culturing of freshly isolated mMG_gl_ or mMG_nb_ with murine GL261 glioblastoma cells. Bottom: Fold-change in colony forming units (CFU) with or without mMG_gl_ or mMG_nb_ co-culturing. (B) Effect of orthotopic co-implantation of mMG_gl_ or mMG_nb_ and GL261 on mice survival. *, p<0.05; **, p<0.0001. n = 10). (C) Effects of co-culturing of GL261 with mMG_gl_ or mMG_nb_ on TMZ resistance, assessed by colony formation assays. *, p<0.05. (D) Ten most abundant proteins identified through unbiased proteomic profiling of conditioned media (CM) derived from immortalized human microglia (hMG). p-value of association between mRNA expression of genes encoding these CM proteins and survival in the TCGA are shown under the TCGA column. To avoid multiple comparisons, only IL11 was tested for survival association using the CGGA. p-value for this association is shown under the CGGA column. (E) Effect of IL11 neutralizing antibodies on hMG CM induced neurosphere formation in SF767. *, p<0.05. ns: not significant. (F) Effect of recombinant human IL11 (20 ng/ml) on neurosphere forming capacity of SF767 cells. *, p<0.05. (G) Survival of mice bearing GL261 cells ectopically expressing IL11 compared to mince baring GL261 cells. n=10. p=0.0003. (H) Effect of ectopic IL11 expression on TMZ resistance of SF767 cells. *In vitro* cell viability was determined. *, p<0.05. n=6. Error bars, SD. (I) GFP-labelled GL261intracranial tumors were sorted into tumor cells (GFP^+^CD45^−^), TAM (GFP-CD45^high^CD11b^+^Gr1^+^), and microglia (GFP-CD45^low^CD11b^+^Gr1^−^). IL11 levels were examined by qRT-PCR, and the results were displayed as heat map. (J) IL11 levels in non-myeloid cells (Iba1^−^CD68^−^CD11b^−^) and tumor associated microglia/myeloid cells (Iba1^+^CD68^+^CD11b^+^) sorted from three glioblastoma specimens resected from human subjects. (K) Effect of microglia co-implantation and ectopic IL11 expression on the expression of the 82 survival-associated genes. RNA was extracted from subcutaneous tumors formed by SF767-vector, SF767-IL11, and SF767 cells co-implanted with microglia and gene expression was analyzed by qRT-PCR. ΔCt between gene and actin in each sample was plotted as heat map. Three independent tumors were analyzed for each cohort.

To determine the potential role of secreted factors in microglia-mediated tumor growth, glioblastoma cells were co-cultured with conditioned medium (CM) derived from hMG (Fig. S4A) or mMG_gl_. These experiments demonstrate that CM recapitulated the microglia-mediated tumor growth (Fig. S4B, S4C). To identify the soluble factor responsible for this phenotype, we performed an unbiased proteomic profiling of CM derived from hMG (58) (Fig. 2D). The list of CM proteins was crossed referenced to their mRNA expression in clinical specimens and survival outcomes. We reasoned that increased mRNA expression for the CM factor underlying the increased tumorigenicity/resistance to DNA damage would be associated with poor clinical survival. With this rationale, we tested the top ten most abundant proteins in the hMG media for clinical survival association. Of those tested, only interleukin 11 (IL11) exhibited the anticipated survival association. In the TCGA dataset, high IL11 mRNA expression was associated with poor survival (Fig. 2D). We then confirmed this association using in the CGGA dataset (Fig. 2D). To avoid multiple comparisons, we tested only IL11 in this confirmatory analysis (Fig. S4D, S4E). Importantly, IL11 mRNA and protein expression were elevated in clinical glioblastoma specimens (Fig. S4E).

Next, we tested whether IL11 is necessary and sufficient for the induction of glioblastoma tumorigenicity and TMZ resistance. Neutralizing antibodies against IL11 suppressed the pro-tumorigenic effect of microglia CM on the SF767, as determined by both neurosphere (Fig. 2E) and colony-forming assays (Fig. S5A). These results were recapitulated using an independent glioblastoma line, LN229 (Fig. S5B). In gain-of-function studies, direct addition of IL11 ligand at concentration found in clinical glioblastoma (Fig. S4E) specimens enhanced neurosphere- and colony-forming potency of SF767 (Fig. 2F and Fig. S5C) and LN229 (Fig. S5D) by ~3-fold. Finally, ectopic expression of human IL11 (hence noted as IL11) in the SF767 glioblastoma line enhanced *in vivo* xenograft formation in a subcutaneous model (Fig. S5E and S5F). In an orthotopic murine glioblastoma model, the median survival of mice bearing GL261 cells ectopically expressing IL11 (designated GL261-IL11) was shortened by ~8 days relative to mice implanted with GL261-vector cells (Fig. 2G and Fig. S5E). Additionally, ectopic IL11 expression decreased the minimal number of implanted cells required for *in vivo* xenograft formation, further supporting the hypothesis that IL11 enhances glioblastoma tumorigenicity (Fig. S5G). Finally, ectopic IL11 expression significantly augmented TMZ resistance of SF767 glioblastoma cells (Fig. 2H).

To determine the major cell source for IL11 in the glioblastoma microenvironment, we sorted tumor and myeloid cells from GL261 tumors. Glioblastomas formed after orthotopic implant of GFP-labelled GL261 cells were sorted into GFP^+^CD45^−^ tumor cells, GFP^−^CD45^low^CD11b^+^Gr1^−^ microglia fraction, and GFP-CD45^high^CD11b^+^Grl^+^ macrophage cells, and then subjected to IL11 qRT-PCR. IL11 mRNA expression was approximately an order of magnitude higher in the microglia fraction relative to the tumor and the macrophage fractions (Fig. 2I). To connect these observations to human glioblastomas, we sorted cell populations from freshly resected clinical glioblastoma samples using Iba1^+^CD68^+^CD11b^+^ as markers for non-tumoral microglia and TAM. Analysis of samples secured from three unrelated patients revealed that IL11 expression was largely found in the Iba1^+^CD68^+^CD11b^+^ fraction (Fig. 2J and Fig. S6), suggesting that the myeloid cell populations were the major cell source of IL11 in the glioblastoma microenvironment.

We next determined if IL11 expression or presence of microglia contributed to increased expression of the 82 survival-associated genes (Table S2) that we collated through our informatics analysis (Fig. 1A). To this end, intracranial glioblastoma xenografts derived from SF767-vector, SF767-IL11, or SF767 cells co-implanted with hMG were isolated and analyzed by qRT-PCR. Supporting our hypothesis, >50% of the 82 genes showed a >2-fold increase in expression when comparing glioblastomas formed by SF767-IL11 to those formed by SF767-vector cells (Fig. 2K, gene list shown in Table S2). Impressively, >90% of the 82 genes showed >2-fold increase when comparing glioblastoma formed after SF767/microglia co-implant relative to those formed by SF767-vector cells. As controls, we showed that ectopic IL11 expression or microglia co-implantation did not systemically affect the expression of a set of randomly selected genes (Fig. S5H).

### IL11 induces glioblastoma STAT3-MYC signaling

In several cancer types, IL11 facilitates tumorigenesis through the activation of the STAT3 pathway (52, 59–62). However, the role of STAT3 in IL11-mediated glioblastoma tumorigenesis remains an open question. IL11 treatment of independent glioblastoma lines induced a time-dependent increase in the accumulation of the active, tyrosine-705 phosphorylated form of STAT3 (p-STAT3), suggesting that IL11 induces STAT3 activation in glioblastoma cells (Fig. 3A). Moreover, inhibitors of STAT3 activation, including Cryptotanshinone (53) and the JAK2 inhibitor, SAR317461 (54), abolished the effect of IL11 on tumor-sphere formation (Fig. 3B), suggesting STAT3 activation is required for the pro-tumorigenic effects of IL11.

**Fig. 3.**
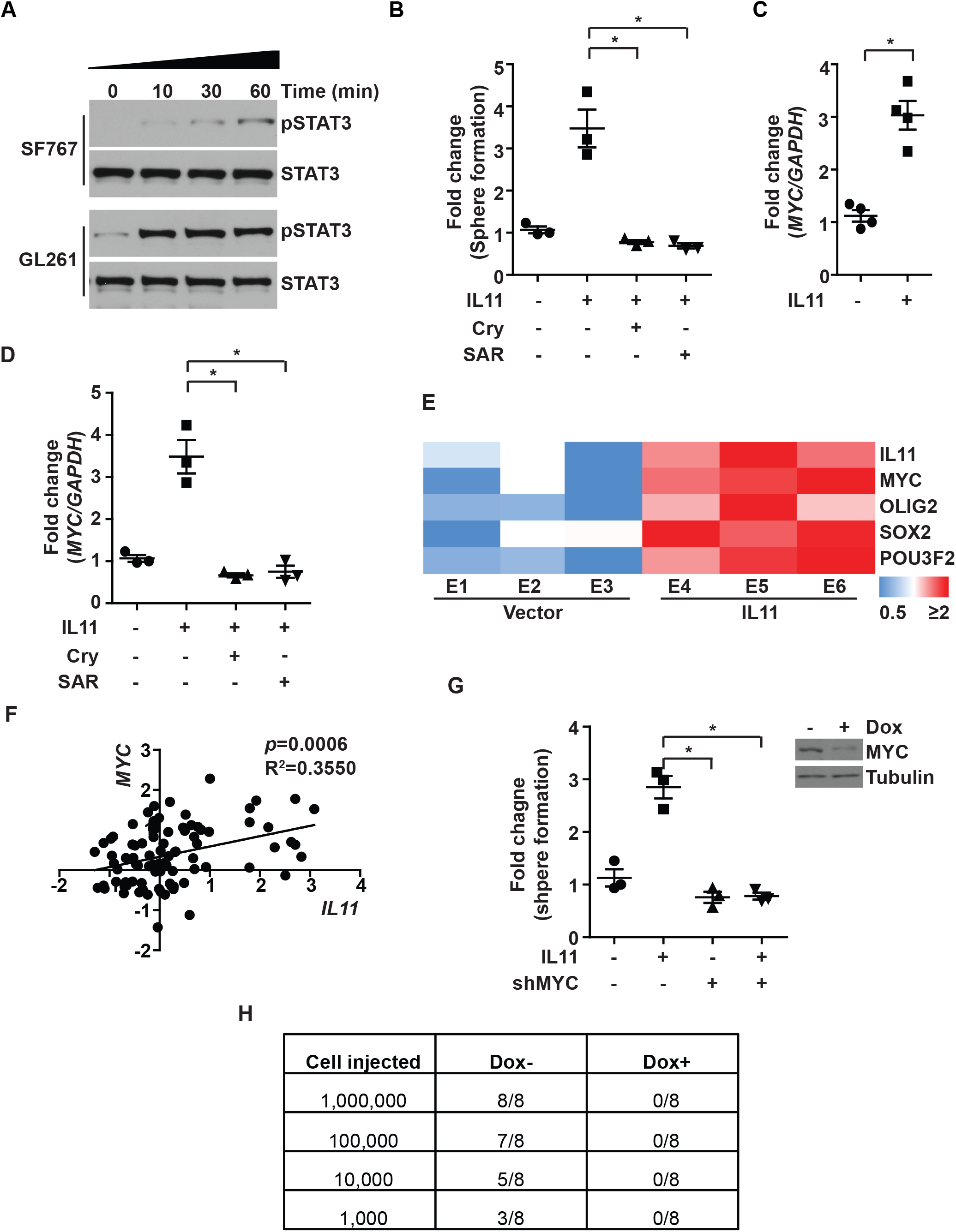
IL11 promotes glioblastoma tumorigenicity via STAT3-MYC signaling. (A) Effects of IL11 stimulation on phospho-STAT3 (Y705) levels in SF767 and GL261 cells. Cells were treated with IL11 (20 ng/ml) for the indicated time prior to immunoblot assay. (B) Effect of IL11 treatment (20 ng/ml) on neurosphere formation in SF767 in the presence or absence of STAT3 inhibitor Cryptotanshinone (Cry) and JAK2 inhibitor SAR317461 (SAR). *, p<0.05. (C) Effect of IL11 treatment (20 ng/ml) on MYC expression, assessed by qRT-PCR. *, p<0.05. (D) Effect of Cryptotanshinone (Cry) and SAR317461 (SAR) treatment on IL11-induced MYC mRNA expression, assessed by qRT-PCR. *, p<0.05. (E) qRT-PCR analysis of MYC, OLIG2, SOX2, and POU3F2 expression in tumors formed after orthotopic implantation of SF767 and SF767 cells that ectopically expressed IL11. ΔCt between gene and actin in each sample was plotted as heat map. Three independent tumors were analyzed for each cohort. (F) Correlation between IL11 (x-axis) and MYC (y-axis) mRNA expression in CGGA glioblastoma patients. R^2^=0.3550, p=0.0006. n=226. (G) Effect of MYC silencing on IL11-induced, *in vitro* neurosphere formation. Dox-inducible shMYC lentivirus was used for stable knockdown of IL11 in SF767 cells. (H) Effect of MYC silencing on IL11 enhanced tumorigenicity *in vivo*. Serial dilutions of SF767-IL11 cells with or without MYC silencing were injected subcutaneously into Nu/Nu mice, and the tumor formation was examined one month after injection. Tumors that grew in size over 50 mm^3^ were scored.

A key downstream effector of STAT3 is MYC (56), a master transcriptional regulator that governs a network of transcriptional factors essential for glioblastoma tumorigenicity, including OLIG2, SOX2, and POU3F2 (63, 64). IL11 treatment of the SF767 glioblastoma line *in vitro* increased *MYC* expression by ~3 fold (Fig. 3C). This increase was abolished by inhibitors that prevent STAT3 activation (Fig. 3D), suggesting an IL11-STAT3-MYC signaling axis. Confirming this observation, the SF767-IL11 xenografts showed increased expression of *MYC* as well as its downstream effectors, *OLIG2*, *SOX2*, and *POU3F2* (63) (Fig. 3E).

Supporting a clinical connection, *IL11* mRNA expression positively correlated with *MYC* mRNA expression in clinical glioblastomas specimens (R^2^=0.3550, *p*=0.0006, CGGA; Fig. 3F). Finally, doxycycline-induced silencing of MYC expression suppressed the effect of IL11 on glioblastoma tumorigenicity *in vitro* (Fig. 3G). For *in vivo* experiments, mice were implanted with serially diluted SF767-IL11 cells harboring a doxycycline-inducible shMYC construct (dox-shMYC) and randomized to doxycycline- or vehicle treatment. By day 30, the vehicle-treated group formed glioblastoma xenografts in a dilution dependent manner (Fig. 3H), while no xenograft was observed in the doxycycline-treated group. Collectively, these results suggest that IL11 in the tumor microenvironment induces activation of a STAT3-MYC axis in glioblastoma cells.

### PI3Kγ inhibition suppresses microglia accumulation and glioblastoma tumorigenicity

PI3Kγ activation in myeloid-derived cells promotes myeloid cell accumulation and re-programming in the tumor microenvironment (65). We hypothesized that PI3Kγ inhibition would impede the accumulation of microglia/macrophages in the glioblastoma microenvironment and effectively convert a poor responder to an exceptional responder. Supporting our hypothesis, treatment of GL261 glioblastoma by the PI3Kγ inhibitor, TG100-115 (65), reduced the density of microglia (CD45^low^CD11b^+^Gr1^−^) by ~2-fold (Fig. 4A). This reduced microglia density was associated with a 3-fold decrease in IL11 concentration in the glioblastoma microenvironment (Fig. 4B). While TG100-115 did not affect glioblastoma growth *in vitro* (Fig. S7A), treatment with TG100-115 prolonged the median survival of GL261 bearing mice by ~5 days relative to mice treated with vehicle (Fig. 4C). Similar survival prolongation was observed when another murine glioblastoma model, CT-2A, was treated with TG100-115 (Fig. 4G).

**Fig. 4.**
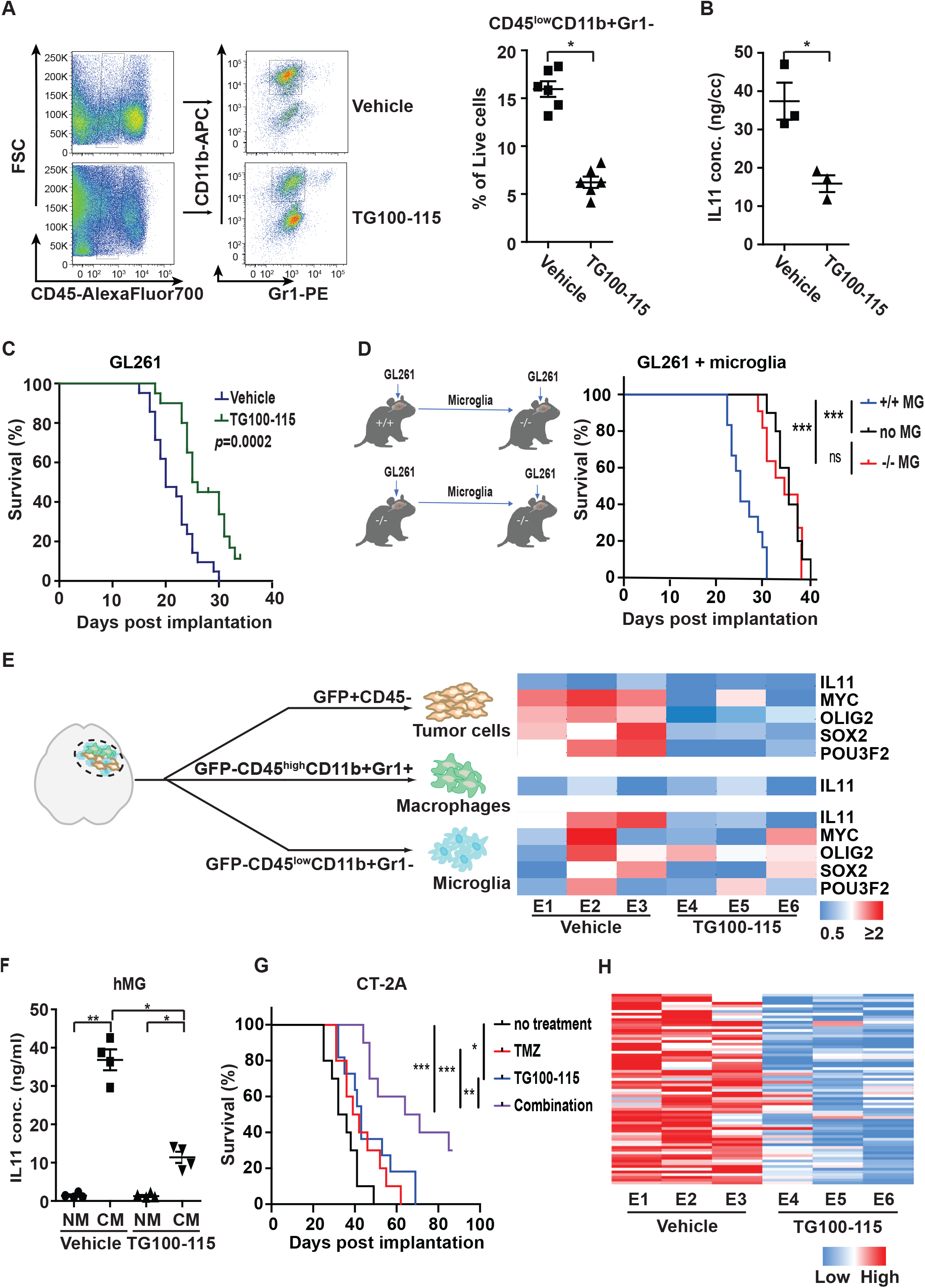
PI3Kγ inhibition suppresses microglia density and glioblastoma tumorigenicity. (A) Effect of treatment with the PI3Kγ inhibitor, TG100-115, on microglia density of tumors formed after orthotopically implant of GL261 cells. Left: Representative flow cytometric analysis. Right: Quantification of microglia density in GL261. n=6; *, p<0.05. (B) ELISA analysis of IL11 protein level in orthotopically implanted GL261 tumors with or without TG100-115 treatment. *, p<0.05. (C) Effect of TG100-115 treatment on survival of mice orthotopically implanted with GL261. n=10, p=0.0002. (D) Survival of PI3Kγ^−/−^ mice after orthotopic co-implant of GL261 cells with the microglia isolated from PI3Kγ^+/+^ or PI3Kγ^−/−^ mice. n=10. ***, p<0.0001; ns: not significant. (E) (Right) qRT-PCR analysis of IL11, MYC, OLIG2, SOX2, and POU3F2 in tumor cells, TAM, and microglia isolated from GL261 tumors that underwent TG100-115 treatment. Orthotopic tumors were resected after treatment and sorted as shown in the left panel. Three independent tumors were analyzed for the treated and untreated cohorts. (F) ELISA analysis of IL levels in CM of TG100-115 treated immortalized human microglia (hMG). (G) Survival of mice orthotopically implanted with CT-2A tumor after treatment with TG100-115, TMZ, or combination of these two drugs (combo). n=10. *, p<0.05; **, p<0.01; ***, p<0.0001. (H) Effect of TG100-115 treatment on the expression of the 82 survival-associated genes. RNA was extracted from orthotopic GL261 tumors with or without TG100-115 treatment and analyzed by qRT-PCR. ΔCt between gene and actin in each sample was plotted as heat map. Three independent tumors were analyzed for the treated and untreated cohorts.

We next confirmed the effects of TG100-115 using genetically engineered mice. *PI3Kγ*^−/−^ C57BL/6 implanted with GL261 glioblastomas exhibited a ~5-day increase in survival relative to GL261 implanted *PI3Kγ*^+/+^ (Fig. S7C). Importantly, TG100-115 treatment of GL261 implanted in *PI3Kγ*^−/−^ mice did not further prolong survival (Fig. S7D). To confirm that PI3Kγ in the microglia population drove this survival difference, microglia (GFP^−^CD45^low^CD11b^+^Gr1^−^) were isolated from glioblastomas formed after orthotopic implant of GL261 cells into *PI3Kγ*^+/+^ and *PI3Kγ*^−/−^ mice. The isolated microglia were then co-implanted with a new batch of GL261 into *PI3Kγ*^−/−^ mice. Consistent with our previous results, there was a ~3-fold reduction in the number of microglia recovered from GL261 implanted *PI3Kγ*^−/−^ mice relative to the *PI3Kγ*^+/+^ mice (Fig. S7E). This reduction was associated with a ~3-fold decrease in IL11 levels (Fig. S7F). Mice implanted with GL261 and microglia isolated from *PI3Kγ*^+/+^ mice exhibited shortened survival by ~ 6 days relative to mice implanted with GL261 and microglia isolated from *PI3Kγ*^−/−^ mice (Fig. 4D). Together, these results support a key role for microglia PI3Kγ in modulating glioblastoma growth.

Next, GFP-labelled GL261 glioblastoma formed with or without TG100-115 treatment were sorted into tumor (GFP^+^CD45^−^), microglia (GFP^−^CD45^low^CD11b^+^Gr1^−^), and macrophages (GFP^−^CD45^high^CD11b^+^Gr1^+^) fractions, and then analyzed for *IL11*, *MYC*, *SOX2*, *OLIG2*, and *POU3F2* mRNA expression. TG100-115 treatment did not significantly alter the already low levels of IL11 expression in the tumor or TAM. However, TG100-115 suppressed microglia IL11 expression by ~3 fold (Fig. 4E). While TG100-115 treatment did not significantly affect the expression of *MYC*, *SOX2*, *OLIG2*, and *POU3F2* in microglia, the expression of these genes in the tumor cells was suppressed by ~2-fold after TG100-115 treatment (Fig. 4E). As these experiments were performed using the same number of sorted cells, the decreased IL11 expression in TG100-115 treated microglia suggested that PI3Kγ inhibition directly suppressed IL11 secretion. Supporting this hypothesis, *in vitro* TG100-115 treatment of cultured hMG suppressed IL11 secretion by ~3 fold (Fig. 4F) without affecting cell viability (Fig. S7G). In aggregate, our results suggest that PI3Kγ inhibition suppressed microglia accumulation and re-programming in the glioblastoma microenvironment.

Supporting our hypothesis that PI3Kγ inhibition transformed the glioblastoma microenvironment from one associated with poor TMZ response to one mimicking that of “exceptional responders”, TG100-115 treatment augmented the anti-neoplastic effects of TMZ. Although TG100-115 treatment failed to affect the TMZ sensitivity of GL261 cells *in vitro* (Fig. S8), substantial efficacy was detected *in vivo*. Survival of mice bearing intracranial CT-2A glioblastomas was prolonged by either TG100115 (~ 7 days) or TMZ treatment (~6 days), but the combination of these treatments extended survival by 31 days (*p*<0.0001, Fig. 4G). Similar results were obtained when this study was carried out using mice bearing intracranial GL261 (Fig. S7B). Further supporting our hypothesis, GL261 tumors treated with TG100-115 (Fig. 4H) or implanted in *PI3Kγ*^−/−^ mice (Fig. S9A) consistently showed decreased expression of the 82 survival-associated gene panel (Table S2). TG100-115 treatment or GL261 engraftment in *PI3Kγ*^−/−^ mice did not systemically affect the expression of a set of randomly selected genes (Fig. S9B). In aggregate, these results suggest that PI3Kγ inhibition converts the glioblastoma tumor microenvironment into one that more closely mimics those observed in “exceptional responders”.

## Discussion

Glioblastomas are diagnosed and graded by pathologic landmarks, including two canonical microenvironmental features, microvascular proliferation and necrosis. Necrotic regions portend a poor prognosis and contain substantial inflammatory components, including microglia and TAM. No consensus exists in terms of the prognostic importance of these cell types (66). The available studies presented conflicting results (67–70) and are largely limited by study designs without considerations for clinical variables that potently influence survival (such as age, KPS, and extent of resection) or validation in independent cohorts. We designed our study to address these limitations. Orthogonal informatics interrogation and cross-validation using three independent patient cohorts unveiled an inverse association between the density of glioblastoma-associated microglia/macrophages and survival. Our results suggest that clinical response to TMZ is, at least in part, modulated by soluble factors released by these glioblastoma-associated microglia/macrophages, including contributions by IL11. To date, IL11’s role in glioblastoma pathogenesis and contribution to the exceptional response remains poorly described. We now show that the microglia/macrophage secreted IL11 activates STAT3-MYC signaling in glioblastoma cells to promote transition into a cancer stem-cell state, resulting in tumorigenicity and TMZ resistance (Fig. 5).

**Fig. 5.**
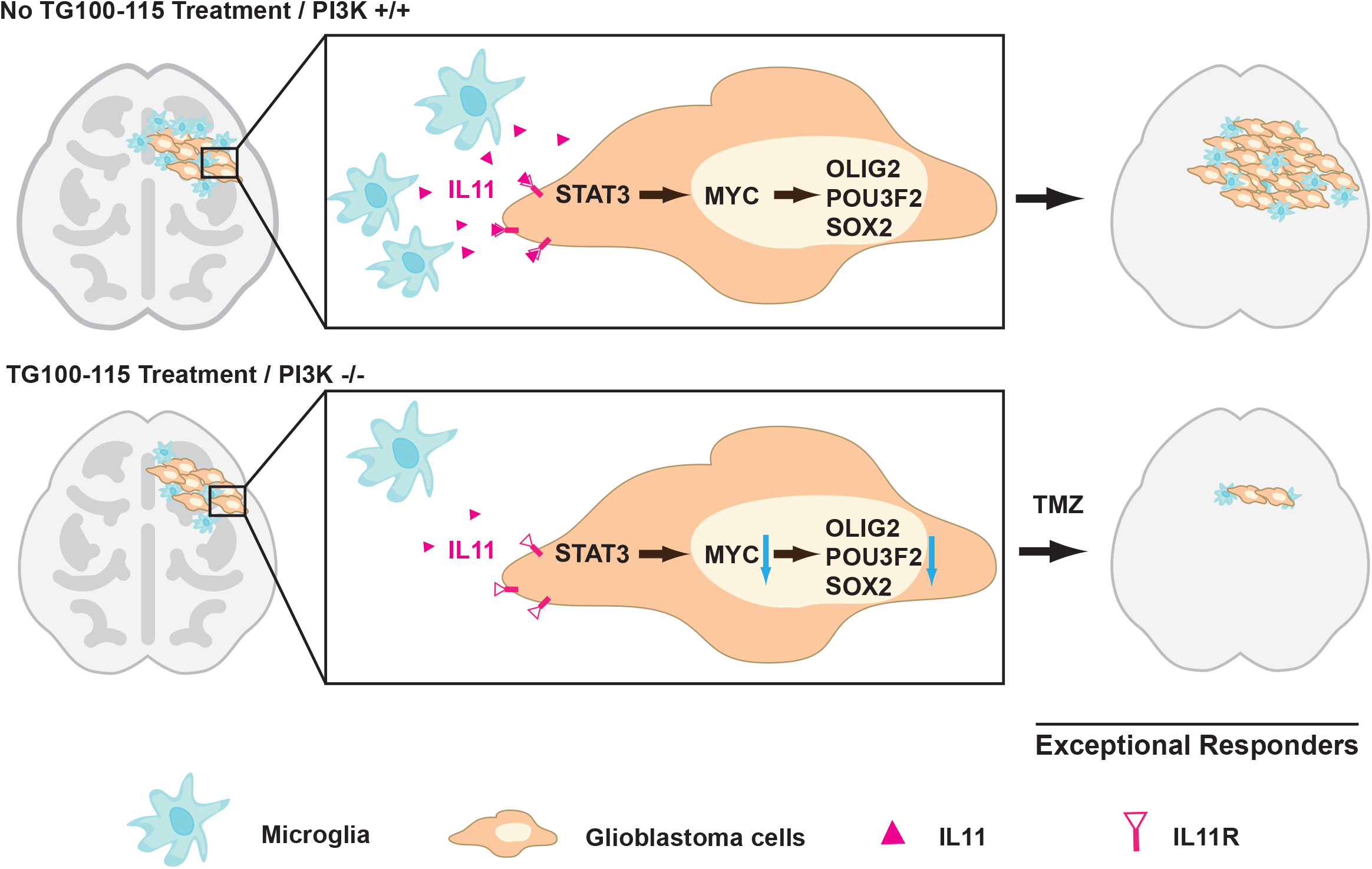
Mechanism underlying “exceptional response” to temozolomide therapy in glioblastoma patients. Glioblastoma-associated microglia secrete high levels of the inflammatory cytokine IL11. The secreted IL11 triggers activation of a STAT3-MYC signaling axis in glioblastoma cells. This signaling induces a change in glioblastoma cell state into a stem-cell like state, causing enhanced tumorigenicity and TMZ resistance. Such signaling is absent in “exceptional responders” and can be suppressed by the PI3Kγ inhibitor, TG100-115 or by PI3Kγ knockout.

Microglia and macrophage in their native state exhibit anti-glioblastoma activities (71). In this context, our findings would suggest that by the time of the clinical presentation, when glioblastoma patients bear significant tumor burden, most microglia/macrophages in the tumor microenvironment have been re-programmed to promote tumor growth (41). The emerging literature on glioblastoma-associated microglia/TAM is largely supportive of this hypothesis (38–40). Further supporting this notion, microglia derived from the normal mice was not pro-tumorigenic (Fig 2B) but became pro-tumorigenic after *in vivo* co-implantation with GL261 glioblastoma cells (Fig. 4E). This re-programming results in altered secretion of a multitude of cytokines, including IL11. Importantly, these cytokines play key roles in inducing peri-tumoral edema that often contributes to clinical deterioration (72). As such, PI3Kγ inhibition has the potential benefit of mitigating deleterious effects resulting from peri-tumoral edema.

We previously demonstrated that stochastic fluctuation in MYC expression influenced glioblastoma cell fate in terms of the equilibrium between entry and exit from the cancer stem cell states (63). Similar findings have been reported for other master-regulatory transcription factors in glioblastomas (64, 73) and for other cancers (74, 75). Our current study suggests that this equilibrium is influenced by glioblastoma microenvironmental factors, including microglia secretion of IL11. Specifically, IL11 activates STAT3 signaling in glioblastoma cells, resulting in increased MYC expression. In turn, increased MYC expression shifts the equilibrium towards the cancer stem cell states. The finding is reminiscent of the observation that post-natal stem cells require signaling from surrounding cells to maintain capacity for self-renewal and multi-potency (76). In this context, re-programming of TAM and microglia may serve as a mechanism by which glioblastoma cells re-constitute signaling required for induction/maintenance of cancer stem cell states.

It remains unclear whether the tumor microenvironment of the exceptional responders is dictated by the intrinsic genetic/epigenetic landscape of the tumor, patient-specific genetics/physiology, or a combination of the two. Recent findings that distinct PIK3CA mutations induce tumor microenvironment with differential capacity for synapse formation suggest the primacy of tumor genetics in shaping its microenvironment (77). Applied to our findings, this model would suggest that microglia/macrophage recruitment is driven by mutations within the cancer genome through cell-autonomous mechanisms. On the other hand, natural polymorphisms in the somatic genome and patient-to-patient variation in neuroendocrine regulation modulate macrophage migration (21) and function (72). As such, aspects of patient physiology independent from tumor genetic/epigenetic composition may contribute to the recruitment and re-programming of microglia/macrophage. As such, non-cell autonomous factors may contribute to microglia/macrophage function in the glioblastoma tumor microenvironment.

Our finding that glioblastoma cells leveraged cellular constituents in its microenvironment to protect itself from the deleterious effects of cancer therapeutics is reminiscent of findings reported in breast and ovarian cancer, where the percentage of stromal cells in the clinical specimen prognosticated survival after systemic therapy (78). These findings bear implications in cancer evolution as it relates to therapeutic resistance (79, 80). Cancer evolution models built on temporal ordering of clonal or sub-clonal mutations typically do not directly account for contributions from the microenvironment (81). While these analyses are of great value in understanding selective pressures imposed on cell-autonomously driven processes, non-cell autonomous phenotypes are invisible to most such analyses. Thus, the trajectory of cancer evolution in response to therapy cannot be fully modeled without understanding the cooperativity of constituents within the complex cancer ecosystem.

In summary, our study suggests clinical glioblastoma specimens derived from exceptional responders to TMZ are characterized by a microenvironment with decreased accumulation of microglia/TAM. This state is recapitulated by PI3Kγ inhibition, which suppresses microglia/macrophage trafficking into glioblastomas. As such, PI3Kγ inhibition warrants consideration in clinical translation as a potential glioblastoma therapeutic.

## Materials and Methods

*Full method details are in SI Appendix, Supplementary Materials and Methods*.

### Literature search and survival signature analysis

The MEDLINE database was queried for studies reporting an mRNA expression signature associated with survival in adult glioblastoma patients. The initial search performed using the following MeSH terms yielded 102 articles: (“prognostic” OR “survival”) AND (“gene signature” OR (“signature” and “gene expression”) OR (“gene signatures”) AND (“glioma” or “glioblastoma” or “GBM”). Two independent reviewers evaluated these articles to identify mRNA signatures associated with glioblastoma survival. 18 signatures were identified (Table S2) and analyzed as described below.

Cohorts of patients with wtIDH were identified in the following manner: IDH status in the TCGA dataset was based on DNA sequencing information; IDH status in the REMBRANDT was assigned based on a published mRNA based gene classifier (46, 82). Single sample gene set enrichment analysis (ssGSEA)(83) score was calculated for each sample for each signature. This enrichment score was normalized to the mean of 10,000 dataset permutations. Cox Proportional Hazard regression was carried out on these normalized enrichment scores to determine survival association.

An mRNA expression dataset of age and KPS matched wtIDH glioblastoma patients was collaged using the Chinese Glioma Genome Atlas (CGGA) with <1-year survival (n=10) and >2-year survival (n=10). Age was matched to ± 5 years, and KPS was precisely matched. ssGSEA enrichment score was calculated for the combined Kim (47) and Lewis (48) signatures for the <1-year and >2-year survivors and compared using t-test. All statistical analyses were carried out in MATLAB (Statistics Toolbox Release 2012b, The MathWorks Inc., Natick, MA). Genes in the signature were compared to a list of 100 genes selected at random from the TCGA. Genes were selected without replacement using the random number generator from R version 3.1. None of the selected genes overlapped with the genes in any of the signatures of interest.

### Cell lines and reagents

Human glioblastoma cell lines LN229 and SF767 were purchased from ATCC. The human microglia (hMG) cell line was purchased from Applied Biological Materials (ABM). Mouse glioblastoma cell GL261 was generously provided by Dr. Santosh Kesari (54). Mouse glioma cell line CT-2A was purchased from EMD Millipore. LN229 and SF767 were cultured in DMEM medium (Gibco) supplemented with 10% fetal bovine serum (FBS, Gibco), 1% Pen-Strep (Gibco) and 1% GlutaMax (Gibco) unless otherwise specified. hMG cells were cultured in Prigrow III medium supplemented with 10% FBS and 1% Pen-Strep. Mouse glioblastoma neurospheres GL261 were cultured in serum-free expansion medium containing DMEM/F12 supplemented with 2% B27 (Invitrogen), 2 μg/ml heparin (StemCell Technologies), epidermal growth factor (EGF, 10 ng/ml; Stem Cell Technology) and basic fibroblast growth factor (bFGF, 20 ng/ml; Stemgent), using ultra-low attachment T-75 flask (Corning). IL11 cDNA was purchased from Origene and cloned into pBabe-puro. shRNA construct shMYC:TRCN0000039642 (Sigma) was cloned into pLKO.1-Tet-Neo obtained from Addgene. For inducible shRNA constructs, transduced cells were treated with or without doxycycline at 1 μg/ml for 5 days before subsequent manipulations. TG100-115 was from Targegen/Sanofi-Aventis (65). TMZ was purchased from AK Scientific.

For collection of conditioned medium (CM), hMG cells were cultured to 20-50×10^6^ cells and resuspended in fresh media for 72 hours. CM were then collected and filtered through a 0.2 μm filter before use in subsequent experiments. IL11 concentration in CM was determined using human and mouse IL11 ELISA kit (Abcam) according to the manufacturer’s protocol. To test the effect of TG100-115 on hMG, cells were treated with 10 μM TG100-115 for 3 days and conditioned medium was collected as described above followed by ELISA assay using human IL11 ELISA kit (Abcam).

### Clinical glioblastoma specimens

Clinical specimen collection was approved by the IRB at the University of California, San Diego Human Research Protections Program (IRB121046) and was in accordance with the principles expressed at the Declaration at Helsinki. The consent process was approved by the ethics committee, and all written consents were documented in our electronic record system.

Pathological analysis of the specimens was performed to verify > 80% tumor content by a board-certified neuropathologist. RNA extraction from clinical glioblastoma specimens was performed using RNeasy mini Kit (Qiagen) according to the manufacturer’s instruction. To determine the levels of IL11 in clinical glioblastoma specimens, specimens were weighed and lysed using RIPA lysis buffer (Sigma) supplemented with a cocktail of protease inhibitors (Roche); IL11 level was then quantitated using the human IL11 ELISA kit (Abcam) according to the manufacturer’s protocol. For experiments involving IL11 FACS, freshly resected glioblastoma samples were dissociated with the Neural Tissue Dissociation Kit (Miltenyi Biotec) followed by myelin removal using Myelin Removal Beads II (Miltenyi Biotec), according to the manufacturer’s instructions. The cell pellet was resuspended in FACS buffer for subsequent fluorescence-activated cell sorting (FACS) isolation.

### Xenograft and allograft models

Animal studies were performed in accordance with the Animal Care and Use Rules at the University of California San Diego under protocol S11253 or University of Minnesota under protocol 1707-349371A. All experiments were conducted in accordance with the “National Institutes of Health Guide for the Care and Use of Laboratory Animals” (NIH publication 80-23). For subcutaneous models, 1×10^6^ of exponentially expanding LN229 or SF767 cells, 1×10^6^ hMG, 0.5×10^6^ SF767 cells mixed with 0.5×10^6^ hMG, or 0.9×10^6^ SF767 mixed with 1×10^5^ mMG_gl_ or mMG_nb_ were implanted in a 50 μl volume into the flanks of Nu/Nu mice (male, ~7 weeks old; Core Facility in University of California, San Diego). To determine the effects of ectopic IL11 expression on glioblastoma growth, SF767 cells transduced with vector or IL11 were implanted as described above. To determine the effect of shMYC on IL11-promoted glioblastoma tumorigenesis, SF767 cells with ectopic IL11 expression were transduced with Dox-inducible shMYC lentivirus, followed by subcutaneous injection as described above. Tumors were counted once tumor reached 50 mm^3^ at 30 days post injection. To facilitate isolation of glioblastoma cells from other cell types in the tumor microenvironment, selective experiments were performed with SF767 cells transduced by a lentiviral construct of GFP-Luc kindly provided by Dr. Dong-Er Zhang (University of California, San Diego). For these experiments, GFP^+^ cells were isolated by FACS 6 weeks post implantation. Tumors were measured with a Vernier caliper twice per week, and tumor volumes were estimated as length × width^2^/2, where Length is the longest diameter and Width is the shortest diameter of the excised tumor. Mice were sacrificed when they developed tumors measuring greater than 2.0 cm in diameter. For experiments involving doxycycline, doxycycline (Sigma) was dissolved at 1 mg/ml in drinking water with 5% sucrose and given to mice starting 7 days after tumor implantation. For experiments involving tumor harvest, SF767 glioblastoma xenografts were resected 30 days after implantation, minced into 0.5 mm pieces, and enzymatically dissociated in PBS containing 0.5mg/ml Collagenase IV (Sigma), 0.1mg/ml Hyaluronidase V (Sigma), 0.6 U/ml Dispase II (Roche) and 0.005 MU/ml DNase I (Sigma) at 37°C for 15 min. The reaction was stopped by washing with PBS, and the cells were centrifuged, washed once with 1x Red Blood Cell lysis buffer (BD) for 5 min at RT, and resuspended in FACS buffer for the subsequent experiments.

For orthotopic co-implantation experiments, dissociated GL261 cells (1×10^6^ cells in 4 μl HBSS), 1×10^6^ GL261 cells mixed with 1×10^5^ mMG_gl_ or mMG_nb_, or CT2A cells (1 × 10^6^ in 4 ul HBBS) were injected into the brains of 6-7 weeks old C57BL/6 mice (male, 6-7 weeks old, The Jackson Laboratory) using murine stereotaxic system (Stoelting Co, Wood Dale, IL). For orthotopic experiments, dissociated GL261 or GL261-GFP/Luc cells (1×10^6^ cells in 4 μl HBSS) were injected into the brains of 6-7 weeks old C57BL/6 mice (The Jackson Laboratory) or *PI3Kγ*^+/+^ or *PI3Kγ*^−/−^ C57BL/6 mice (kindly provided by Dr. Varner at University of California, San Diego). The coordinates were: 1.8 mm to the right of bregma and 3 mm deep from the dura. For experiments involving TG100-115, treatments were initiated 7 days after the initial tumor injection. TG100-115 (2.5 mg/Kg) was administered through i.p. injection every 12 hours for 2-3 weeks.

To discriminate GL261 cells from other cell types in the tumor microenvironment, select experiments were performed using GL261 transduced with a lentiviral construct of GFP-Luc. For experiment involving temozolomide (TMZ), TMZ (50 mg/kg) was administered i.p. daily for 3 cycles as described previously(84). TMZ was dissolved in 5 % DMSO and 5 % solutol-15 in sterile saline. Mouse weight was monitored daily starting 7 days post injection. Mice with >20% reductions in body weight were perfused with PBS and the brains were processed further as specified. Tumor-bearing mice were euthanized by i.p. injection of 250 μl ketamine and xylazine mixture and perfused using PBS. The brain was collected and stored in ice-cold PBS and tumor-bearing regions were resected for further analysis. These samples were processed as above described (under **clinical glioblastoma specimens)**. To measure IL11 level in mouse GL261 orthotopic tumor samples, tumor tissues were resected 21 days post implantation and lysed as described above. The lysates were used to determine IL11 level using mouse IL11 ELISA kit (Abcam).

### Statistical analysis

The CGGA dataset was kindly provided by Dr. Tao Jiang as normalized, probe-level expression values. Values of probes designed to assess the same gene were averaged. Data from the Cancer Genome Atlas (TCGA) data was downloaded from the TCGA data portal (https://tcga-data.nci.nih.gov/tcga/) as level 3 data. Expression levels of ten most abundant cytokines in the conditioned media hMG were identified by proteomic profiling. To validate the survival association of IL11, the TCGA data set was divided into two groups using a median cut off point dependent on IL11 expression level, for which a Kaplan-Meier survival curve was plotted. Significance was tested by unpaired two-tailed Student’s t-test using SPSS. This analysis was exploratory and did not use a multi-comparison correction. The association between IL11 and survival was subsequently validated in an independent dataset (CGGA) using the same method and Kaplan Meier survival curve was plotted. For the association between MYC and IL11, expression levels of MYC and IL11 in CGGA data set were plotted at Y axis and X axis, separately. Data are presented as means with their respective standard error of the mean (SEM) or as statistical scatter plots generated using GraphPad Prism 5. For survival curves related to GL261 and CT-2A allograft studies, p-values were analyzed using the log-rank (Mantel-Cox) test. *P*<0.05 was considered to be statistically significant. Student’s t-test (two-tailed) was used in other statistical comparisons unless otherwise indicated above.

## Acknowledgements

The work is supported by NIH grants NS097649-01 and CA240953-01, the Doris Duke Charitable Foundation Clinical Scientist Development Award (www.ddcf.org), the Sontag Foundation Distinguished Scientist Award (www.sontagfoundation.org), the Burroughs Wellcome Fund Career Awards for Medical Scientists (www.bwfund.org), the Kimmel Scholar award (www.kimmel.org), and a Discovery Grant from the American Brain Tumor Association (www.abta.org). We thank Dr. Bing Ren for helpful comments. We are grateful to Brian Hirshman, Valya Ramakrishnan, and Johnny Akers for their assistance in qRT-PCR, informatics analysis, animal care, and graphical design.

## Author contributions

Flow cytometry studies were performed by JL, MMK, and JM. *In vivo* animal studies and analyses were performed by JL, MMK, JN, HZ, GY, SD, TK, and BX. *In vitro* studies were performed by JL, and JM. Informatic analysis was performed by KP, AS, TJ, GY. All studies were conceived, directed and analyzed by ML, FF, TJ, AW, CKG, JAV and CCC. The manuscript was written by JL, ASV, JNR, JM, and CCC.

