## Supplemental Figures and Tables for "PI3Kγ inhibition suppresses microglia/TAM accumulation in glioblastoma microenvironment to promote exceptional temozolomide response"

**This PDF file includes:**

SI Materials and Methods
Figs. S1 to S9
Tables S1 to S3

**Supplementary Information Materials and Methods.**

**Colony formation, neurosphere formation, and viability assays**

For colony formation assays, LN229 ($0.5\times10^3$ cells/well) or SF767 ($3\times10^3$ cells/well) cells were seeded in 6-well plates and treated as indicated. Colonies were fixed, stained with 0.5% crystal violet in methanol, and scored after 14 days. For neurosphere formation assays, LN229, SF767 or GL261 single cells were seeded into individual wells of ultra-low-attachment 96-well plates (Corning) containing 100 µl neurosphere media followed by treatment as indicated. Wells were scored after 10-14 days for aggregates of ≥ 50 cells. Approximately 100 wells were scored in each experiment.

To test the effect of TG100-115 on the viability in immortalized hMG or GL261 glioblastoma cells, cells were plated at $2\times10^3$/well in 96-well plates and treated with TG100-115 at the indicated concentrations for 3 days. Cell viability was measured using CellTiter-Blu cell viability assay kit (Promega).

**Immunoblotting**

Lysates were prepared using a RIPA lysis buffer (Sigma) supplemented with a cocktail of protease inhibitors (Roche), incubated for 15 min on ice, and then clarified by centrifugation. Protein lysates were quantitated using Bicinchoninic acid (BCA) assay. Equal amounts of protein were fractionated using SDS-polyacrylamide gel electrophoresis and electro-transferred to nitrocellulose membranes (Invitrogen). Membranes were blocked for 1 h in 5% fat-free milk dissolved in TBS containing 0.1% Tween-20 and incubated overnight at 4°C with indicated antibodies, including anti-HA for IL11(Cell Signaling Technology), anti-phospho-STAT3 (Tyr-705) (Cell Signaling Technology), and total STAT3 (Cell Signaling Technology). For loading control, membranes were probed with anti-tubulin (Sigma). After washing, membranes were incubated with appropriate secondary (Pierce) antibodies conjugated to horseradish peroxidase (Pierce).

**Flow cytometry staining, analysis, and sorting**

Single cell suspensions ($10^6$ cells in 100 µl total volume) were incubated with Aqua Live Dead fixable stain at 4°C for 15 min (Life Technology), FcR-blocking antibody at 4°C for 10 min (BD

Biosciences), and various fluorescently labeled antibodies at 4°C for 1h. Primary antibodies against CD45-Alexa700 (mouse, 30-F11), CD11b-APC (mouse, M1/70), Gr1-FITC (mouse, RB6-8C5), CD3e-eFluor780 (mouse, 145-2C11), CD68-PE (human, eBioY1/82A), CD11b-APC (human, ICRF44) were purchased from eBioscience; F4/80-PE-Cy7 (mouse, BM8), CD4-PEdazzle (mouse, GK1.5), CD8-BV605 (mouse, 53-13) were purchased from Biolegend. Iba1-FITC (human) was from Abcam. For intracellular staining, cells were fixed and permeabilized using Flow Cytometry Permeabilization/Wash Buffer I (R&D) and then incubated with rabbit IL11 antibody (R&D) followed by PE-Cy7 conjugated anti-Rabbit secondary antibody (eBioscience), or cells were permeabilized using Permeabilization/Wash Buffer I (R&D) and then incubated with anti-Iba1-FITC antibody (Abcam). Multicolor FACS analysis was performed on a BD Canto RUO19 Color Analyzer at Flow Cytometry Core at the UC San Diego Center for AIDS Research. All data analysis was performed using FlowJo (Treestar).
For cell sorting, single cell suspensions were stained with Aqua Live Dead fixable stain (Life Technologies) to exclude dead cells and anti-CD11b-APC (mouse, M1/70, eBioscience), anti-Gr1-PE (RB6-8C5, eBioscience). FACS sorting was performed using FACS Aria 11 color highspeed sorter at the Flow Cytometry Core at UC San Diego Center for AIDS Research. GFP-tagged SF767 tumor cells were sorted based on GFP positivity. GFP-tagged GL261 cells were sorted into $GFP^{+}CD45^{-}$ (glioblastoma), $GFP^{-}CD45^{low}CD11b^{+}Gr1^{-}$ (microglia), and GFP-$CD45^{high}CD11b^{+}Gr1^{+}$ (macrophages). Microglia were isolated from both GL261 glioblastomas ($mMG_{gl}$) and normal murine brain ($mMG_{nb}$). Cells from freshly resected glioblastoma specimens were sorted into Iba1-CD68-CD11b- (non-myeloid cells) and $Iba1^{+}CD68^{+}CD11b^{+}$ (microglia/myeloid cells).

**Supplemental Figure legends**

**Fig. S1. GCESS based comparison of TCGA GBM dataset to REMBRANDT GBM Dataset.** Parallel survival associated gene clusters are identified in both the TCGA_u133 (A) and REMBRANDT (B) dataset. Arrows represent cluster 2 in both datasets associated with enrichment of macrophage/microglia gene signatures. TCGA: n=539. REMBRANDT: n=227. (C) REMBRANDT cluster 2 is associated with poor outcome of GBM patients. High and low expression were defined based on the median expression value. p=0.0325.

**Fig. S2. hMG promotes glioblastoma tumorigenicity and TMZ resistance.** (A) Effect of Co-culturing of immortalized human microglia (hMG) with SF767 (left) or LN229 (right) glioblastoma cells on colony formation potency. The result was displayed as fold-change in colony forming unit (CFU) with or without co-culturing. -, tumor cells only; +, co-culturing with indicated cell types. (B) Effect of co-injection of hMG and SF767 on tumor growth in mice. Tumor growth was monitored twice per week and mean tumor volume±SD are shown. n=6; *, p<0.05. Bottom: Representative images of H&E staining of glioblastoma formed. Scale bar, 50 µm. (C) Effect of hMG co-culturing with SF767 on TMZ resistance detected by colony formation assay. *, p<0.05. (D) Survival of mice with intracranial co-implantation of freshly isolated glioblastoma-associated macrophages mMPgl ($GFP^{-}CD45^{high}CD11b^{+}Gr1^{+}$) and GL261. n=10-12. p<0.0001.

**Fig. S3. Gating strategy for flow cytometric analysis of myeloid cells and microglia in glioblastoma specimens.** Cells from freshly resected glioblastoma specimens were sorted into $CD45^{+}CD11b^{+}GR1^{+}$ (myeloid cells) and $CD45^{-}CD11b^{+}Gr1^{-}$ (microglia).

**Fig. S4. Identification of IL11 in the CM isolated from hMG.** (A) Effect of hMG-derived conditioned medium (CM) on colony (left) and sphere formation (right) in SF767 cells. Left: Representative images of colony formation assay were shown. Fold change in colony forming unit (CFU) with or without co-culturing was labeled. Right: Quantification of sphere formation assay. *, p<0.05. (B) Effect of conditioned medium (CM) isolated from $mMG_{gl}$ on colony formation of GL261. *, p<0.01. (C) Effect of $mMG_{gl}$ co-injection with SF767 on tumor xenograft growth. Tumor growth was monitored twice per week and mean tumor volume±SD are shown. n=6; *, p<0.05. (D) Kaplan-Meier analysis of the association of IL11 expression levels and GBM patient survival. CGGA glioblastoma patients were defined as high or low IL11 expressing based on median cut-off. p=0.019. n=226. (E) Upper: Elevated IL11 mRNA expression in glioblastoma specimens relative to surrounding normal brain tissues was detected by qRT-PCR. T, Tumor; N, Tumor adjacent normal tissue. Bottom: ELISA analysis showed elevated IL11 levels in glioblastoma specimens relative to surrounding brain tissues. *, p<0.05; **, p<0.01.

**Fig. S5. Microglia/myeloid secreted IL11 enhances glioblastoma tumorigenicity and TMZ resistance.** (A) Effect of IL11 with or without neutralizing antibodies treatment on SF767 colony formation potency. *, p<0.05. (B) Effect of IL11 with or without nneutralizing antibodies treatment on LN229 colony (left) and sphere (right) formation potency. *, p<0.05. (C) Treatment with recombinant IL11 augmented colony forming potency in SF767 cells. *, p<0.05. (D) Effect of recombinant IL11 on colony (left) and sphere (right) forming potency in LN229 cells. *, p<0.05. (E) Ectopic expression of IL11 in SF767 cells (left) and GL261 (right) was examined by qRT-PCR. *, p<0.05. (F) Effect of ectopic expression of human IL11 on SF767 subcutaneous tumor growth. *n*=6. Error bars, SD. (G) Effect of ectopic expression of human IL11 on SF767 *in vivo* tumorigenicity. Serial dilutions of SF767 cells with or without human IL11 overexpression were injected subcutaneously into Nu/Nu mice, and the tumor formation was examined one month after injection. Tumors that grew in size over 50 $mm^3$ were counted. (H) IL11 expression did not alter the expression level of genes selected using a random number generator. RNA was extracted from subcutaneous tumors formed by SF767-vector, SF767-IL11 and SF767 co-implanted with hMG and gene expression was analyzed by qRT-PCR. ΔCt between gene and actin in each sample was plotted as heat map.

**Fig. S6. Gating strategy for flow cytometric analysis of non-myeloid cells and microglia/myeloid in clinical glioblastoma specimens.** Cells from freshly resected glioblastoma specimens were sorted into $Iba1^-CD68^-CD11b^-$ (non myeloid cells) and $Iba1^+CD68^+CD11b^+$ (microglia/myeloid cells) and stained for IL11 expression.

**Fig. S7. The effect of genetic or pharmacological PI3Kγ inhibition on myeloid accumulation and glioblastoma tumorigenicity.** (A) Effect of TG100-115 treatment on the viability of GL261 cells *in vitro*. (B) Effect of TG100-115, TMZ, and combination treatment on mice survival implanted with GL261. n=10. p<0.001. (C) Comparing survival of PI3Kγ$^{-/-}$ and PI3Kγ$^{+/+}$ C57BL/6 mice implanted with GL261 glioblastomas. p<0.001. n=10. (D) Effect of TG100-115 treatment on PI3K$\gamma^{+/+}$ and *PI3K*$\gamma^{-/-}$ mice implanted with GL261 cells. n=10. ns, not significant. (E) Left: Flow cytometric analysis of microglia (CD45$^{low}$CD11b$^{+}$Gr1$^{-}$) in GL261 orthotopic tumors implanted in *PI3K*$\gamma^{+/+}$ and *PI3K*$\gamma^{-/-}$ C57BL/6 mice. Right: Quantification of microglia population in GL261 intracranial murine model. *n*=6; *, p<0.05. (F) Decreased IL11 level in the GL261 orthotopical mouse model implanted in *PI3K*$\gamma^{-/-}$ mice compared to those in *PI3K*$\gamma^{+/+}$ mice. IL11 protein levels were analyzed by ELISA. *, p < 0.05. (G) Effect of TG100-115 treatment on the viability of hMG cells *in vitro*. ns, not significant.

**Fig. S8. TG100-115 treatment fails to affect the TMZ sensitivity of GL261 cells *in vitro.*** SF767 cells were treated with escalating doses of TG100-15 and TMZ. Cell viability was measured using CellTiter-Blu cell viability assay kit.

**Fig. S9. The effect of genetic or pharmacological PI3Kγ inhibition on the expression of clinical survival-associated signatures or randomly selected genes.** (A) Effect of genetic inhibition of PI3Kγ on the expression of the survival-associated inflammatory signatures was assessed by qRT-PCR. (B) The expression levels of the randomly selected genes affected by pharmacological (left) or genetic (right) inhibition of PI3Kγ were analyzed by qRT-PCR. ΔCt between gene and actin in each sample was plotted as heat map.

Fig S_1

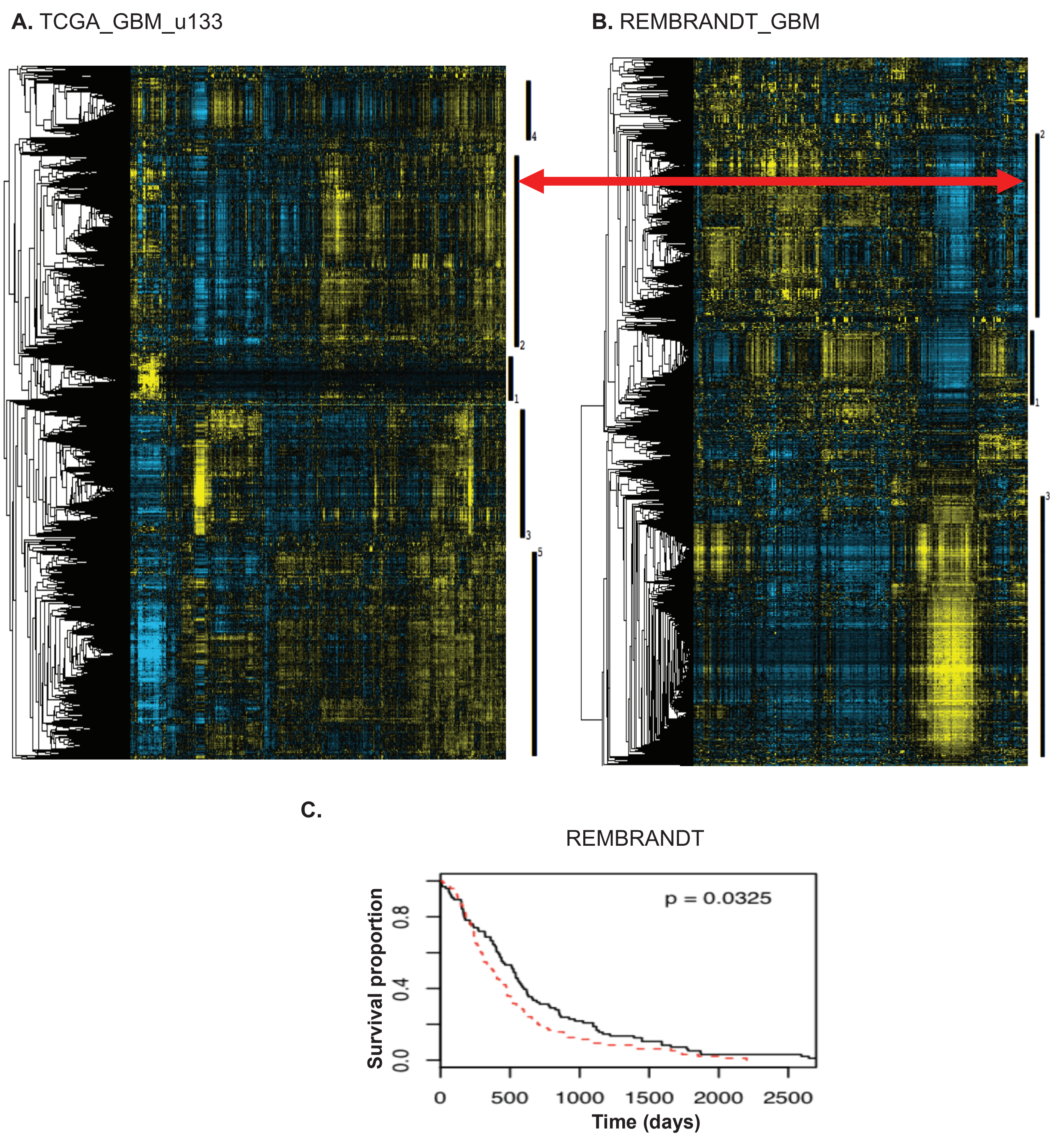
A. TCGA_GBM_u133
4
2
1
3
5
B. REMBRANDT_GBM
2
1
3
C.
REMBRANDT
p = 0.0325
Survival proportion
0.0
0.4
0.8
0
500
1000
1500
2000
2500
Time (days)

Fig S_2

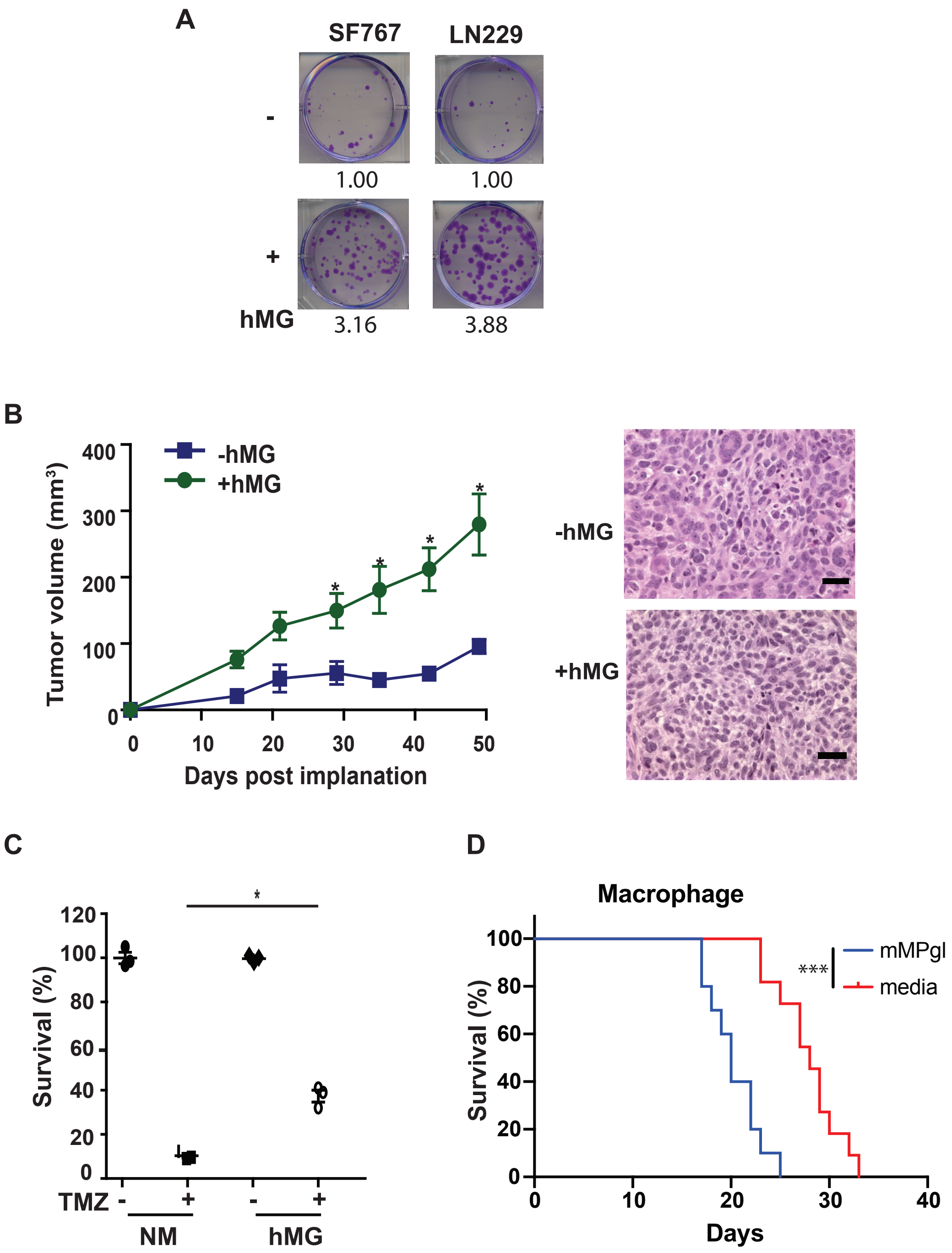
A
SF767
LN229
-
1.00
1.00
+
hMG
3.16
3.88
B
Tumor volume (mm³)
400
300
200
100
0
-hMG
+hMG
*
0
10
20
30
40
50
Days post implanation
-hMG
+hMG
C
Survival (%)
120
100
80
60
40
20
0
TMZ
-
+
-
+
NM
hMG
D
Macrophage
Survival (%)
100
80
60
40
20
0
***
mMPgl
media
0
10
20
30
40
Days

Fig S_3

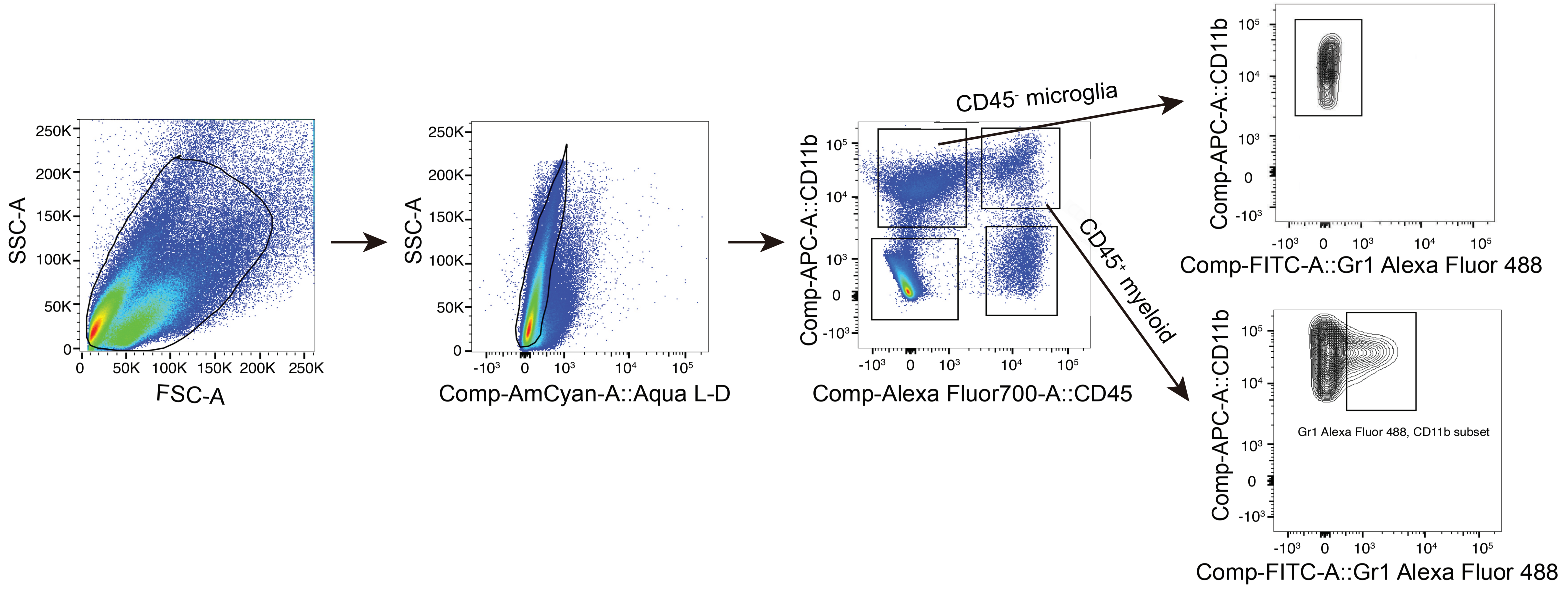

SSC-A
FSC-A
Comp-AmCyan-A::Aqua L-D
Comp-APC-A::CD11b
Comp-Alexa Fluor700-A::CD45
CD45- microglia
CD45+ myeloid
Comp-FITC-A::Gr1 Alexa Fluor 488
Gr1 Alexa Fluor 488, CD11b subset

Fig S_4

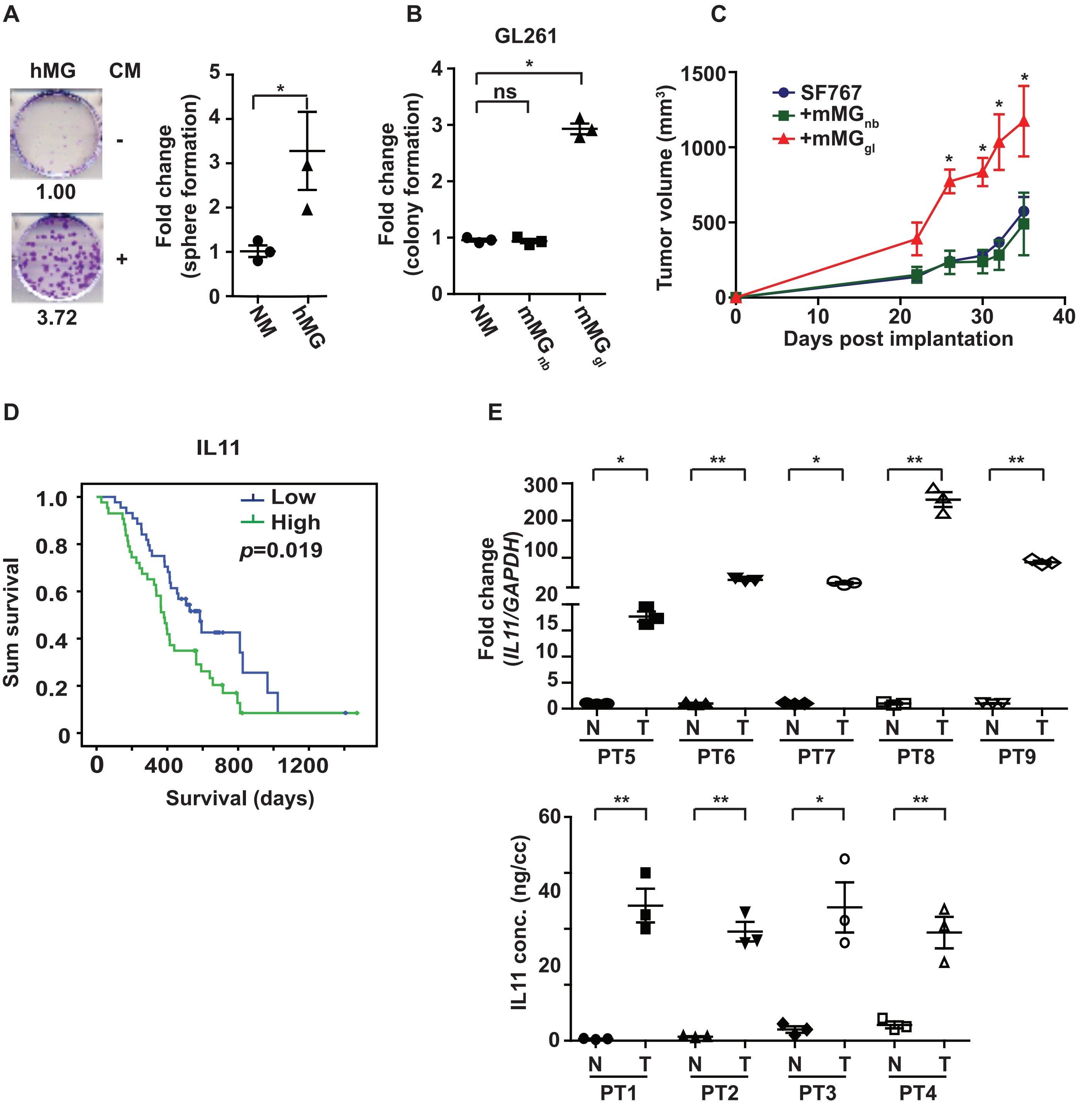
A
hMG
CM
-
+
1.00
3.72
Fold change (sphere formation)
NM
hMG
B
GL261
ns
Fold change (colony formation)
NM
mMG nb
mMG gl
C
SF767
+mMG nb
+mMG gl
Tumor volume (mm3)
Days post implantation
D
IL11
Low
High
p=0.019
Sum survival
Survival (days)
E
Fold change (IL11/GAPDH)
N T
PT5
PT6
PT7
PT8
PT9
IL11 conc. (ng/cc)
PT1
PT2
PT3
PT4

Fig S_5

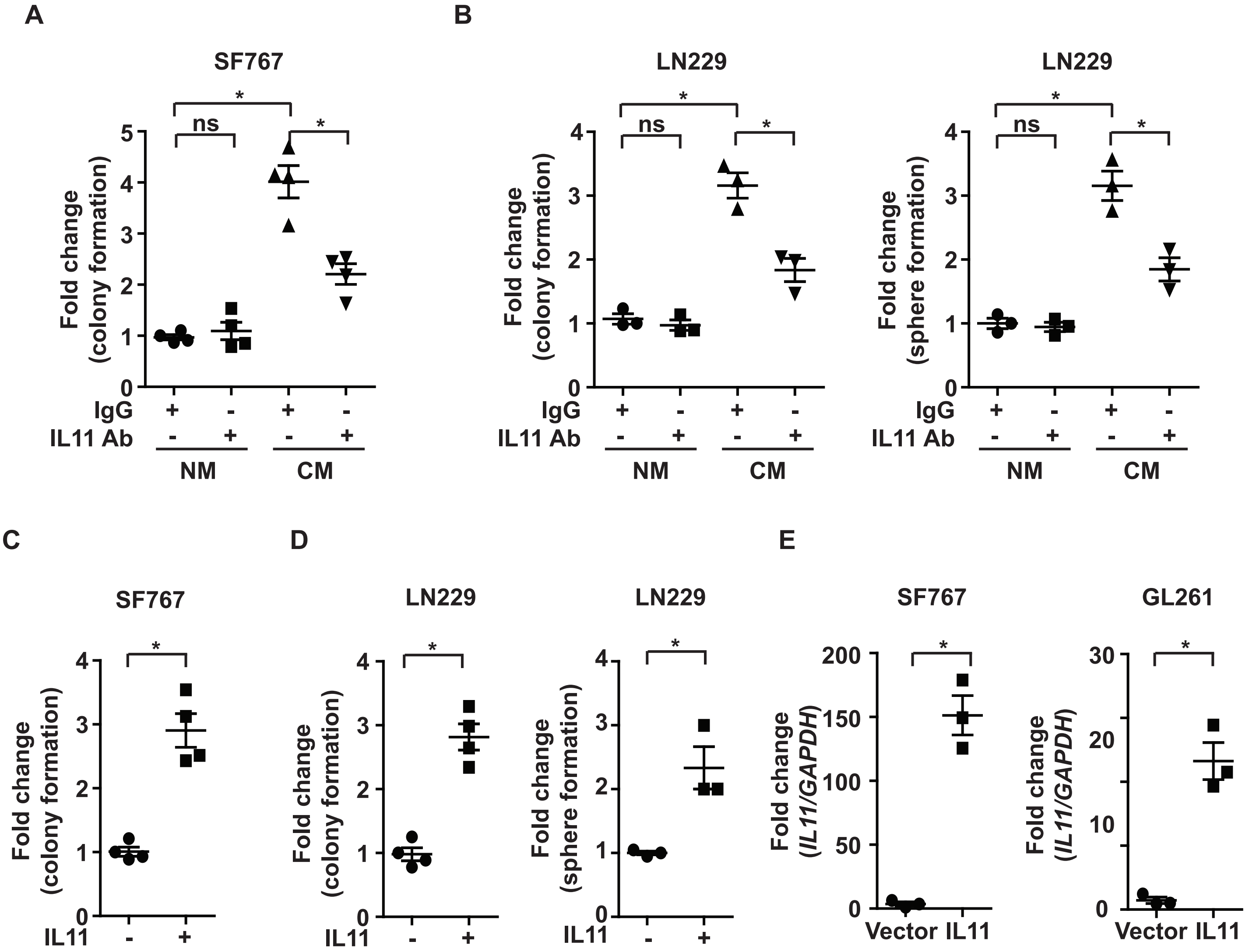


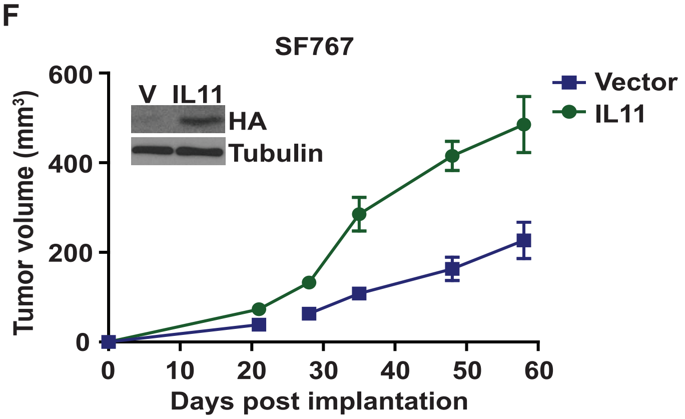


G

| Cell injected | Vector | IL11 |
|---|---|---|
| 1,000,000 | 8/8 | 8/8 |
| 100,000 | 6/8 | 8/8 |
| 10,000 | 2/8 | 8/8 |
| 1,000 | 0/8 | 4/8 |

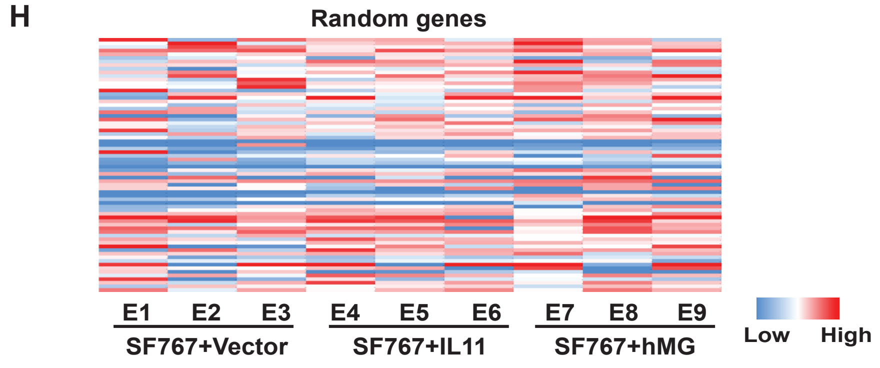

### Fig S_6

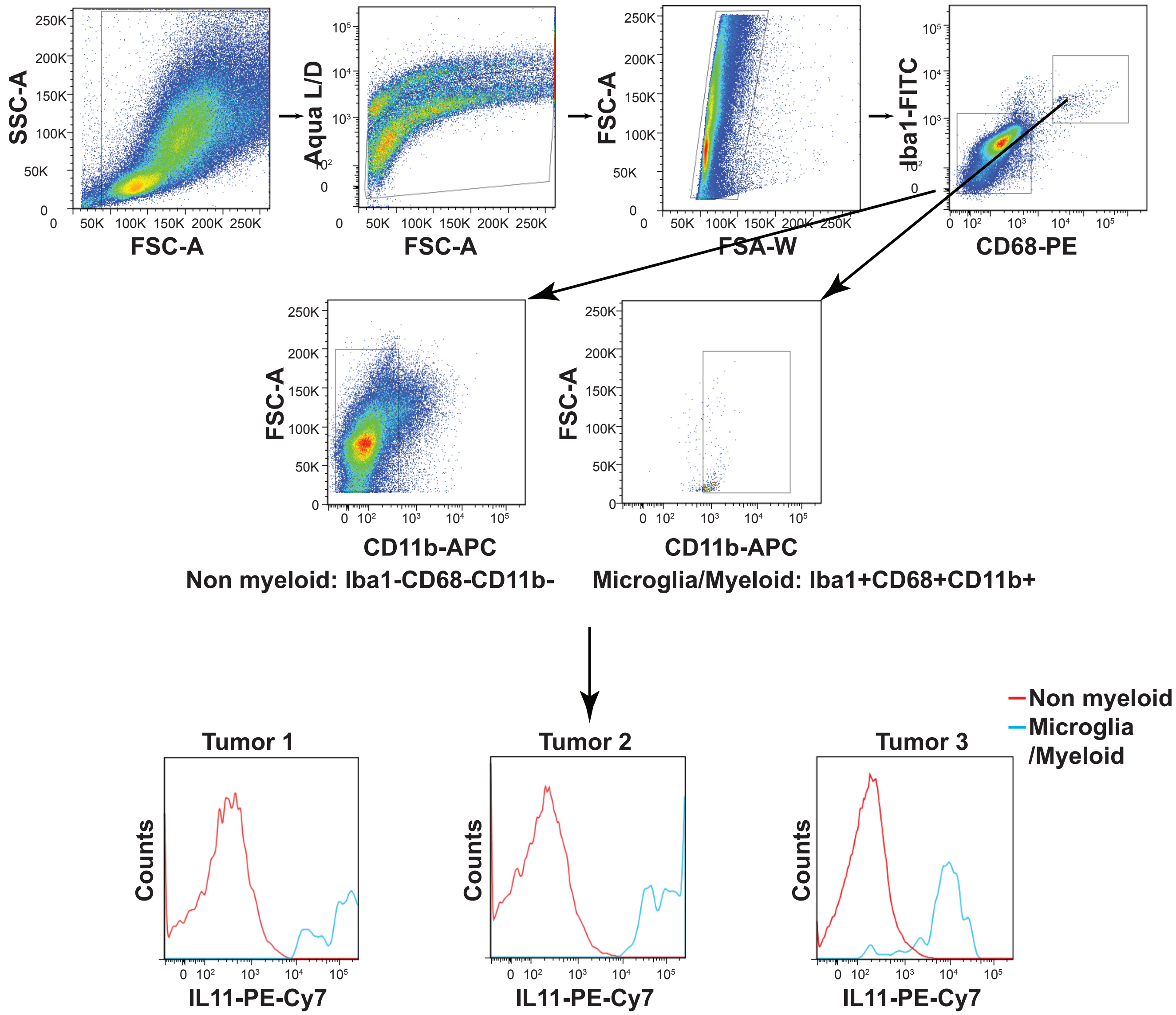

SSC-A
FSC-A
Aqua L/D
FSC-A
FSA-W
Iba1-FITC
CD68-PE
CD11b-APC
Non myeloid: Iba1-CD68-CD11b-
CD11b-APC
Microglia/Myeloid: Iba1+CD68+CD11b+
Non myeloid
Microglia /Myeloid
Tumor 1
Tumor 2
Tumor 3
Counts
IL11-PE-Cy7

Fig S_7

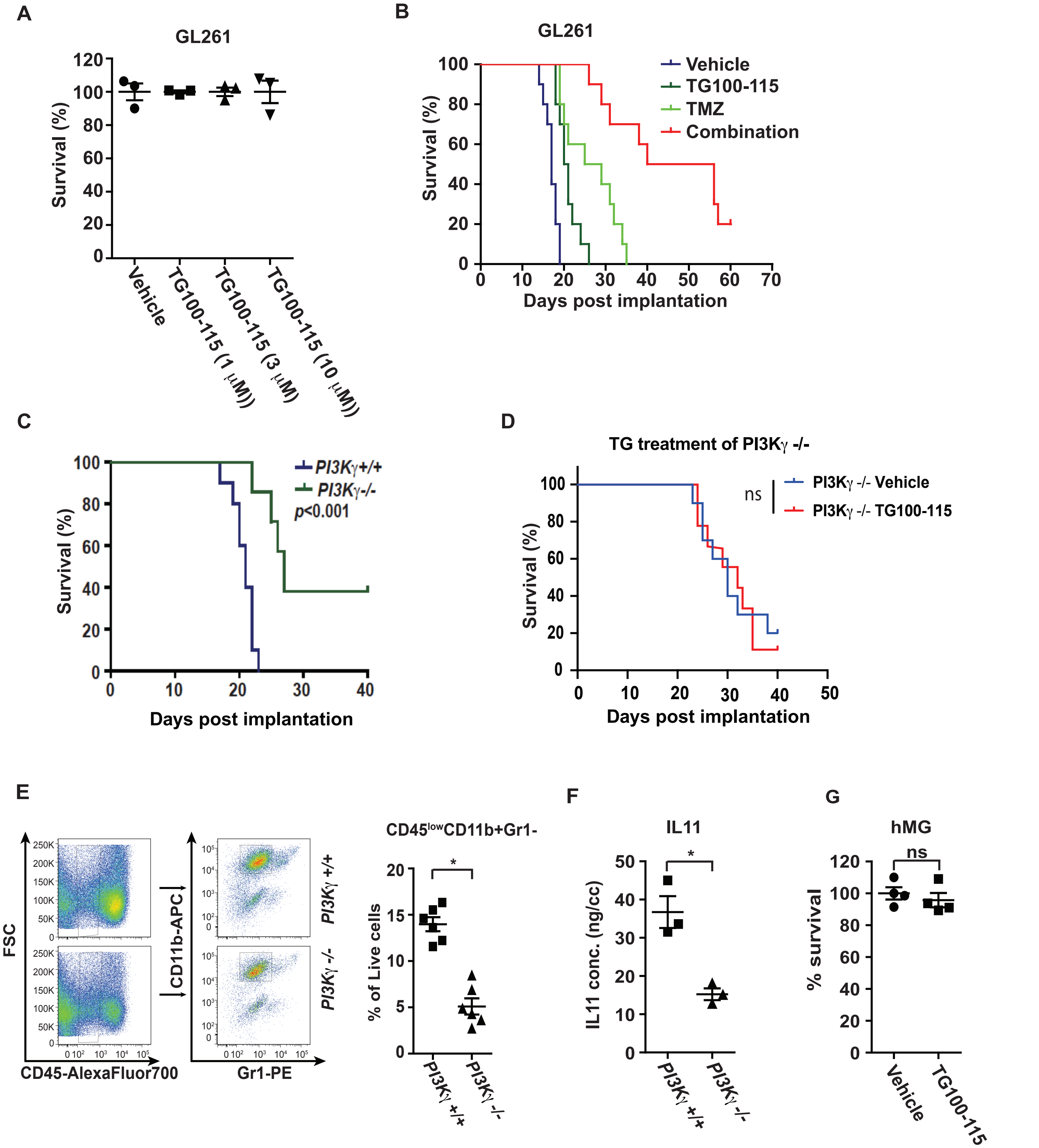
A
GL261
Survival (%)
Vehicle
TG100-115 (1 μM)
TG100-115 (3 μM)
TG100-115 (10 μM)
B
GL261
Vehicle
TG100-115
TMZ
Combination
Survival (%)
Days post implantation
C
PI3Kγ+/+
PI3Kγ-/-
p<0.001
Survival (%)
Days post implantation
D
TG treatment of PI3Kγ -/-
ns
PI3Kγ -/- Vehicle
PI3Kγ -/- TG100-115
Survival (%)
Days post implantation
E
FSC
CD45-AlexaFluor700
CD11b-APC
Gr1-PE
PI3Kγ +/+
PI3Kγ -/-
CD45lowCD11b+Gr1-
% of Live cells
F
IL11
IL11 conc. (ng/cc)
PI3Kγ +/+
PI3Kγ -/-
G
hMG
ns
% survival
Vehicle
TG100-115

**Fig S_8**

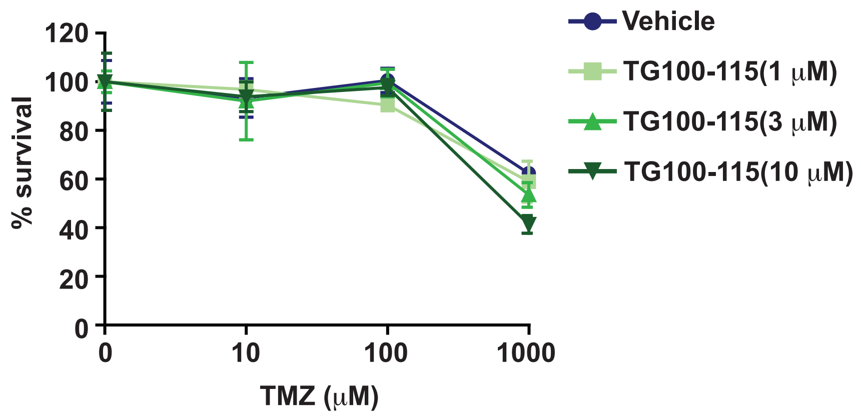

Vehicle
TG100-115(1 μM)
TG100-115(3 μM)
TG100-115(10 μM)
% survival
120
100
80
60
40
20
0
0
10
100
1000
TMZ (μM)

Fig S_9

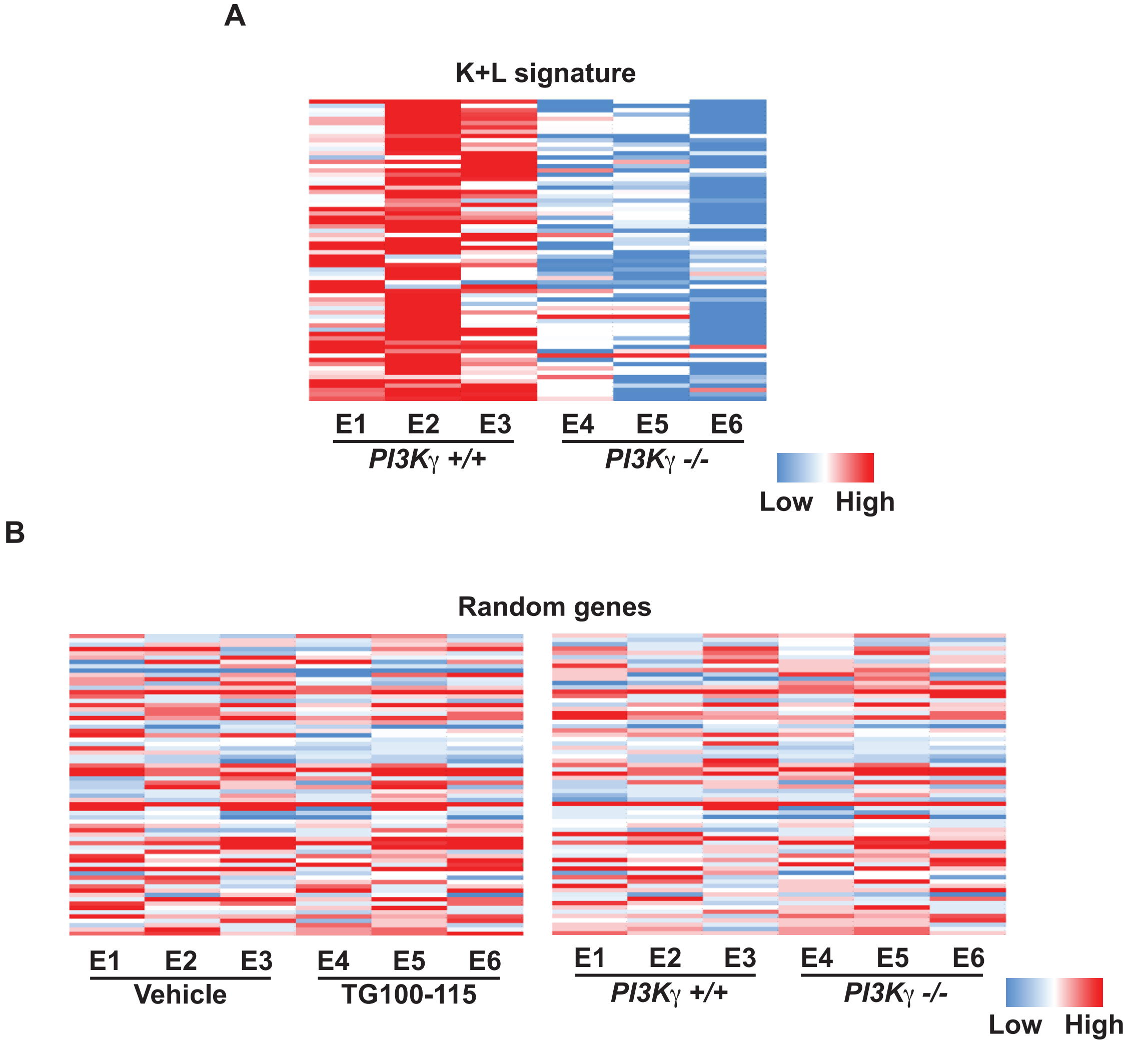
A
K+L signature
E1 E2 E3
PI3Kγ +/+
E4 E5 E6
PI3Kγ -/-
Low High
B
Random genes
E1 E2 E3
Vehicle
E4 E5 E6
TG100-115
E1 E2 E3
PI3Kγ +/+
E4 E5 E6
PI3Kγ -/-
Low High

**Table S1: Kim and Lewis glioblastoma survival gene signatures showed survival association in TCGA and REMBRANDT wtIDH cohorts (*: *p*<0.05).**

| Signature | PubMed ID | wtIDH | |
|---|---|---|---|
| | | TCGA | REM |
| Arimappamagan (2013) | 23646114 | * | |
| Bao (2013) | 24279471 | | |
| Bredel (2006) | 16365179 | | |
| Duarte (2012) | 22242177 | | |
| Huang (2014) | 25537983 | * | |
| Kawaguchi (2011) | 22021018 | * | |
| Kim (2013) | 23502430 | * | * |
| Lewis (2015) | 25619842 | * | * |
| Li (2015) | 25628952 | | |
| Meng (2014) | 24970813 | | |
| Mikheev (2015) | 25140038 | | |
| Murat (2008) | 18565887 | | * |
| Ng (2012) | 22675171 | | |
| Noushmehr (2010) | 20399149 | | * |
| Reifenberger (2014) | 24615357 | | * |
| Shukla (2013) | 24078801 | | * |
| de Tayrac (2011) | 21224364 | | * |
| Zhou (2014) | 24714365 | | |

**Table S2. The list of genes in Kim and Lewis glioblastoma survival-associated signatures.**

| Kim | | Lewis | |
|---|---|---|---|
| DEPDC6 | LGALS8 | PMP22 | NUPR1 |
| RPRM | DYNLT3 | FABP7 | GPNMB |
| NET1 | KIAA0323 | SERPINA3 | IL1A |
| WAC | TFRC | FAM134B | NR1H3 |
| March8 | KIAA0495 | GPR56 | APOBEC3F |
| AI054381 | FBX017 | CAP2 | HFE |
| REPS2 | TMEM22 | PLEKHA4 | PPCS |
| ZNF609 | LOC3900940 | SLC22A4 | S100A13 |
| KLF13 | MT1E | SLC22A5 | P4HA2 |
| IL8 | DCTD | CSRP1 | SLC20A1 |
| ADM | FLJ11286 | CHMP2A | CXADR |
| PDPN | C13orf18 | PICK1 | MDK |
| IGFBP2 | HOMER1 | BEST1 | RBP1 |
| MDK | FAM3C | VGF | C10orf10 |
| TIMP1 | CASP3 | SEZ6L2 | CFI |
| EFEMP2 | NSUN5 | ASPHD1 | IFI30 |
| ACOX2 | PDLIM3 | DBNDD1 | SERPINA1 |
| TAGLN2 | MT1M | SMPD1 | C5AR1 |
| SLC43A3 | | ALDOA | TNFRSF1B |
| | | CYCS | CD14 |
| | | RNF14 | RNASET2 |
| | | C7orf25 | GLB1 |

**Table S3: The list of qRT-PCR primers.**

| Gene | Forward | Reverse |
|---|---|---|
| DEPDC6 | 5'-ATAGACGGCACCATCTCAAAAC-3' | 5'-GTCGGCTAATTTCTGCATGAGT-3' |
| RPRM | 5'-GTGGTGCAGATCGCAGTCAT-3' | 5'-CGGTCCTTCACTAGGAAGTTGA-3' |
| Net1 | 5'-GTGGCGCATGATGAGATCG-3' | 5'-CATCTAAGACTCGGATCGTCCTT-3' |
| MAC | 5'-TCAGTGATGGCTGTCACGAC-3' | 5'-TTTGGTGGTGAAGGATCTGCG-3' |
| MARCH8 | 5'-TTCTCAGGATGCCATTTCTGC-3' | 5'-GTTGCTTGGATGACTCATGGAA-3' |
| AI054381 | 5'-CTACGCCGCCATCAATGATTC-3' | 5'-GGGATAGTGGTAGGGCTTCAC-3' |
| REPS2 | 5'-GATGGTGAGATACGGTTTGGG-3' | 5'-TCAAGGACACCCTCTGGTAATG-3' |
| ZNF609 | 5'-TCCAAAAGGAGTAAGAGTGGCA- | 5'-CTACACTGCGTGACGCCTTAG-3' |
| KLF13 | 5'-CCTGGCCTCAGACAAAGGG-3' | 5'-ATTTCCCGTAAACTTTCTCGCA-3' |
| CXCL15-IL8 | 5'-CAAGGCTGGTCCATGCTCC-3' | 5'-TGCTATCACTTCCTTTCTGTTGC-3' |
| ADM | 5'-CACCCTGATGTTATTGGGTTCA-3' | 5'-TTAGCGCCCACTTATTCCACT-3' |
| PDPN | 5'-ACCGTGCCAGTGTTGTTCTG-3' | 5'-AGCACCTGTGGTTGTTATTTTGT-3' |
| IGFBP2 | 5'-CAGACGCTACGCTGCTATCC-3' | 5'-CCCTCAGAGTGGTCGTCATCA-3' |
| MDK | 5'-GAAGAAGGCGCGGTACAATG-3' | 5'-GAGGTGCAGGGCTTAGTCA-3' |
| TIMP1 | 5'-GCAACTCGGACCTGGTCATAA-3' | 5'-CGGCCCGTGATGAGAAACT-3' |
| EFEMP2 | 5'-CGGACAGCTACACGGAATG-3' | 5'-CGAGGCAGACACAAATAACCC-3' |
| ACOX2 | 5'-ACGGTCCTGAACGCATTTATG-3' | 5'-TTGGCCCCATTTAGCAATCTG-3' |
| TAGLN2 | 5'-CCTGGCCGTGAGAACTTCC-3' | 5'-GTCCGTGGTGTTAATGCCATAG-3' |
| SLC43A3 | 5'-CCTTGTTGACTGGACTCTTGG-3' | 5'-AGGCCCTGTTACATTGCTCTG-3' |
| LGALS8 | 5'-TTCAAAAGGTCTAGCTGCATTGT- | 5'-CCATGAACACGATCTCAAAGGA-3' |
| DYNLT3 | 5'-TGGACTGCAAGCATAGTGGAA-3' | 5'-GTGAAATCCATACGGGCTCCT-3' |
| KIAA0323- | 5'-GAGCCCACGGTTTCTTCCTT-3' | 5'-CCATCACAAACGCCTCGGTTA-3' |
| FBXO17 | 5'-GCTGCCGACAAGTCTCTCAC-3' | 5'-GAGTTACAAGGGCACCGTAGT-3' |
| TMEM22 | 5'-CTTGTCCGTGATAGTTGTGTGT-3' | 5'-AGGTATAAGCACAGGTGATGGA-3' |
| MT1E | 5'-TCAGGTTGGGAGGGAACTCAA-3' | 5'-GAAAGCCTGGAGAGGGAATGA-3' |
| DCTD | 5'-ATGGCCCGAGTATTTCATGGC-3' | 5'-CATCACTGCACCCATTTGGC-3' |
| FLJ 11286 | 5'-CTCCATCGTGTACGGGGTAAA-3' | 5'-GGTCCTGCTTCATATCTTCTGGT-3' |
| C13orf18- | 5'-TTCCTGGATCAACCCGCTG-3' | 5'-CTGTAGGACAATATCGAAAGGCG- |
| HOMER1 | 5'-CTTCACAGGAATCAGCAGGAG-3' | 5'-GTCCCATTGATACTTTCTGGTGT-3' |
| FAM3C | 5'-GGACTCAGCCATTCGTTCTAC-3' | 5'-GCTGCTCCACTAGCCATCTTAAA-3' |
| CASP3 | 5'-TGGTGATGAAGGGGTCATTTATG- | 5'-TTCGGCTTTCCAGTCAGACTC-3' |
| NSUN5 | 5'-CTGAAGCAGTTGTACGCTCTG-3' | 5'-CCCTTCCCCAGCAATAATTCAT-3' |
| PDLM3 | 5'-TGGGGGCATAGACTTCAATCA-3' | 5'-CTCCGTACCAAAGCCATCAATAG-3' |
| PMP22 | 5'-CATCGCGGTGCTAGTGTTG-3' | 5'-AAGGCGGATGTGGTACAGTTC-3' |
| FABP7 | 5'-GGACACAATGCACATTCAAGAAC- | 5'-CCGAACCACAGACTTACAGTTT-3' |
| SERPINA3 | 5'-ATTTGTCCCAATGTCTGCGAA-3' | 5'-TGGCTATCTTGGCTATAAAGGGG-3' |
| FAM134B | 5'-AAACAGCAGAGTCCTGGCAAG-3' | 5'-AGGTAGCTGAGTATGACCCCA-3' |
| GRP56- | 5'-CTGCGGCAGATGGTCTACTTC-3' | 5'-ATAGTGGAGGGTGCTCTGTTG-3' |
| CAP2 | 5'-CAGGGTTAGACGGACCTCC-3' | 5'-CTCGGCCACCATACTGTTTATC-3' |
| PLEXHA4 | 5'-AGCACCAAGAAGCCTACCC-3' | 5'-CGTCGAAGTGCATTGTTCCTTT-3' |
| SLC22A4 | 5'-TGGTATGTCAGTCGTGTTCCT-3' | 5'-AGCCCCATCGCAGAGAAGT-3' |
| SLC22A5 | 5'-ACTGTGCCAGGGGTGCTAT-3' | 5'-GCAACTGAGGCTTCGTAGAAT-3' |

| | | |
|---|---|---|
| CSRP1 | 5'-AACAGCTTCCATAAATCCTGCTT- | 5'-CCATACTTCTTGCCGTAACATGA-3' |
| CHMP2A | 5'-AGACGCCAGAGGAACTACTTC-3' | 5'-ACCAGGTCTTTTGCCATGATTC-3' |
| PICK1 | 5'-CGCTAGGCGAGCCCTATAC-3' | 5'-CTTGGACATGGTGGATACGAAG-3' |
| BEST1 | 5'-ACACAACACATTCTGGGTGC-3' | 5'-CGCAAAGTACACACCTCATTCA-3' |
| VGF | 5'-AAGGATGACGGCGTACCAGA-3' | 5'-TGCCTGCAACAGTACCGAG-3' |
| SEZ6L2 | 5'-GGAGGATGAGATGATGCCAGA-3' | 5'-GGTGGCTAGTGTGGGGTCT-3' |
| ASPHD1 | 5'-CCACCAATGCTCGGGTTAGAT-3' | 5'-CGAAGACAAAGTCGAGAGCTT-3' |
| DBNDD1 | 5'-CTGCCTACGTTCACCATCCTG-3' | 5'-GGCTGTTTCTCACGGTTCTG-3' |
| SMPD1 | 5'-TGGGACTCCTTTGGATGGG-3' | 5'-CGGCGCTATGGCACTGAAT-3' |
| ALDOA | 5'-CGTGTGAATCCCTGCATTGG-3' | 5'-CAGCCCCTGGGTAGTTGTC-3' |
| CYCS | 5'-CCAAATCTCCACGGTCTGTTC-3' | 5'-ATCAGGGTATCCTCTCCCCAG-3' |
| RNF14 | 5'-TACTGGCCCTGGCGAGTATTT-3' | 5'-CCTTGGACAGACTCAGCTTTC-3' |
| C7ORF25- | 5'-CATGCTCTCCGACAGAATTGT-3' | 5'-CGATTCCACGATGGCTTTCAG-3' |
| ARF4 | 5'-CTGGCAAGACGACAATTCTGT-3' | 5'-CCACAAAAATGAGACCCTGGGTA- |
| NUPR1 | 5'-CCCTTCCCAGCAACCTCTAAA-3' | 5'-TCTTGGTCCGACCTTTCCGA-3' |
| GPNMB | 5'-AGAAATGGAGCTTTGTCTACGTC- | 5'-CTTCGAGATGGGAATGTATGCC-3' |
| IL1A | 5'-GCACCTTACACCTACCAGAGT-3' | 5'-AAACTTCTGCCTGACGAGCTT-3' |
| NR1H3 | 5'-ACAGAGCTTCGTCCACAAAAG-3' | 5'-GCGTGCTCCCTTGATGACA-3' |
| APOBEC3F | 5'-TTTCTCGTCGGAATACCGTCT-3' | 5'-CAAAGGGGCTACTGAGCACTT-3' |
| HFE | 5'-CACCGCGTTCACATTCTCTAA-3' | 5'-CTGGCTTGAGGTTTGCTCC-3' |
| PPCS | 5'-CGCTTTCTGGACAACTTCAGT-3' | 5'-GGGAGCGCATTCTCTTCGG-3' |
| S100A13 | 5'-AACTGCCTCATTTGCTCAAGG-3' | 5'-AGTCTCCAGTATTCACTGAACCT-3' |
| P4HA2 | 5'-CAGGTACTATGATGTGATGTCCG- | 5'-AAAGGGTCGCCTTGAGAAGTC-3' |
| SLC20A1 | 5'-TTTGACAAACTTCCTCTGTGGG-3' | 5'-GGACTTTCAGACGGACTAGACTT-3' |
| CXADR | 5'-GCACCCGCTAAGGTAGCTG-3' | 5'-ATAGACCCGTCCTTGCTCTGT-3' |
| MDK | 5'-CGCGGTCGCCAAAAAGAAAG-3' | 5'-TACTTGCAGTCGGCTCCAAAC-3' |
| RBP1 | 5'-CTGAGCAATGAGAATTTCGAGGA- | 5'-GCGGTCGTCTATGCCTGTC-3' |
| C10ORF10 | 5'-TGCCCACAATTCGGGAGAC-3' | 5'-AGACCTCACGTAGTCATCCAG-3' |
| CFI | 5'-CTTGGCTCTCCACTTGAGTTC-3' | 5'-GGAGCGATGCGTGTATTTCTG-3' |
| IFI30 | 5'-CCTGGTCTCCGATCCTACCAT-3' | 5'-TTGCAGGTGGTTGTGCCTT-3' |
| SERPINA1 | 5'-GAGCATTGGCACAGCGTTTG-3' | 5'-AAGCGATGGTTGGATGTCAGC-3' |
| C5AR1 | 5'-TACCATTAGTGCCGACCGTTT-3' | 5'-CCGGTACACGAAGGATGGAAT-3' |
| TNFRSF1B | 5'-ACACCCTACAAACCGGAACC-3' | 5'-AGCCTTCCTGTCATAGTATTCCT-3' |
| CD14 | 5'-CTCTGTCCTTAAAGCGGCTTAC-3' | 5'-GTTGCGGAGGTTCAAGATGTT-3' |
| RNASET2A | 5'-CCCCCAACAGTATGCAAGGAG-3' | 5'-AAGCTAGTTTTCCCAGACTGC-3' |
| GLB1 | 5'-GCACGGCATCTATAATGTCACC-3' | 5'-GTATCGGAATGGCTGTCCATC-3' |
| DAPK3 | 5'-GCACGACATCTTCGAGAACAA-3' | 5'-CTTAGAGTGCAGGTAGTGAACG-3' |
| RANBP9 | 5'-GGCCGTGGACGAACAAGAG-3' | 5'-GAACTGACGCGGCATCTTTT-3' |
| ACY1 | 5'-GGCACCAACCCTACACTCTC-3' | 5'-ACTCCAATGTTCCTTGAAGACAG-3' |
| BRF1 | 5'-GGTGTGCCTCACGCATCTC-3' | 5'-GAAGGCCCCTCATCCTACAGA-3' |
| APOBEC1 | 5'-TGAGGAGAAGAATCGAACCCT-3' | 5'-CTTGATTTCGTAGAGCAGACAGG-3' |
| LRFN3 | 5'-ACACTGGAGGACCTCGACC-3' | 5'-CTTGTGCAGGCGGGAAAAAG-3' |
| SOCS7 | 5'-GGGTCAAGACAGTCGGTGG-3' | 5'-TCTCGGCCTCCGATTCCAA-3' |
| VBP1 | 5'-AGTCCACCAACTCAATGGAGA-3' | 5'-CAAGCATTACATTAGCCCCCAA-3' |

| | | |
|---|---|---|
| IRS4 | 5'-CGACCAAGCGACAAGAAGACT-3' | 5'-GGTTCCCGAGGAAAGAAGCG-3' |
| DEFB4 | 5'-GGTGGTATAGGCGATCCTGTT-3' | 5'-AGGGCAAAAGACTGGATGACA-3' |
| FOXL2 | 5'-GGTCGCACAGTCAAGGAGC-3' | 5'-CGCGATGATGTACTGGTAGATG-3' |
| CENPN | 5'-TGAACTGACAACAATCCTGAAGG- | 5'-CTTGCACGCTTTTCCTCACAC-3' |
| ITGB3BP | 5'-AATCTCCTGTGCATCACATTTCT-3' | 5'-TTCATAGCTGTCAAGATGACGTG-3' |
| SLCO1B3 | 5'-TGGAGCAACAGTACGGTCAG-3' | 5'-TGCTTTCGCAGATTAGAGGGAA-3' |
| PRPS2 | 5'-AGCTCGCATCAGGACCTGT-3' | 5'-ACGCTTTCACCAATCTCCACG-3' |
| PLSCR3 | 5'-TGTGGCCTCCAGGAGATGGA-3' | 5'-GGAGGAAGGGATGCCAGGTC-3' |
| S100A10 | 5'-GGCTACTTAACAAAGGAGGACC-3' | 5'-GAGGCCCGCAATTAGGGAAA-3' |
| FAM110B | 5'-TAGCTCCGAGGGCTCTAGC-3' | 5'-CACCTTGCGGATGTCCGAA-3' |
| CCDC59 | 5'-CGGCCTGGTGGTATTGAGG-3' | 5'-GCCGCCATGTCTTCTGTCT-3' |
| HLF | 5'-CCACCTTTATCCCGCCTCC-3' | 5'-TTTACTAAATGCGTCTTCGTGGT-3' |
| LAMC1 | 5'-GGACTCCGCCCGAGGAATA-3' | 5'-ACTTGAGACGCACATAGGTGA-3' |
| CORIN | 5'-CCAAAGCCGGTCTTGAGAG-3' | 5'-GAGGAGGTTAGCAGTCGCC-3' |
| FAM130A2 | 5'-AAAAGGGAGAAACGACTGAGGA- | 5'-TGGGCACACTTGTGAAGCC-3' |
| ANKZF1 | 5'-ATGCTCCGGTCTTTCAGGG-3' | 5'-GGTCTGGTCACAAGTTGAACAAA- |
| DNM1 | 5'-ATATGCCGAGTTCCTGCACTG-3' | 5'-AGTAGACGCGGAGGTTGATAG-3' |
| METTL8 | 5'-TGCACTGTCCTACTGTGCCT-3' | 5'-TCCAGCTCCACAACCAACCT-3' |
| STK17A | 5'-TCTGAGTCGGCTGTTGATTTC-3' | 5'-GGGGTGCTTTAGACATTCTTCA-3' |
| C17orf63 | 5'-CAGCCGAATTGGATGCGTATG-3' | 5'-TCACAGTACGACGAACGTGTT-3' |
| ME1 | 5'-CTGCTGACACGGAACCCTC-3' | 5'-GATCTCCTGACTGTTGAAGGAAG-3' |
| GTF2B | 5'-ACTACAGAGCCGGTGATATGAT-3' | 5'-GTTGCTTTGTCATTGCTGAAAGT-3' |
| FDXR | 5'-CTGAGGCAGAGTCGAGTGAAG-3' | 5'-CCCGAAGCTCCTTAATGGTGA-3' |
| KIAA0888 | 5'-GTGGATATGCTGGAAAATTGCAG- | 5'-CAGGGTCCCCACACCTTAAAT-3' |
| DGCR2 | 5'-GGTGGCACCACTACGAAGG-3' | 5'-AGGTGGCGAGAGAGCCATT-3' |
| DHRS9 | 5'-CTGTGGACTCGTAAAGGAAAACT- | 5'-GCAGCGATTACATGAAATCCCT-3' |
| RPGRIP1L | 5'-ATGTCTGGTCCAACTGATGAGA-3' | 5'-TCCGTGTTGTTGAAGTTTCTTGT-3' |
| HSPA6 | 5'-CAAGGTGCGCGTATGCTAC-3' | 5'-GCTCATTGATGATCCGCAACAC-3' |
| TRA@ | 5'-TGCCGTGTACCAGCTGAGAG-3' | 5'-GCCACAGCACTGTTGCTCTT-3' |
| IGSF6 | 5'-ATGAGGCCGTCACCATAAAGT-3' | 5'-GCACCGTAGCGAAACCACA-3' |
| DNAJB5 | 5'-AAGATGGCCTTGAAGTACCACC-3' | 5'-GCCTCTGCAATCTCCTTAAACT-3' |
| LUC7L | 5'-GGACGCGCATGGATTTAGGA-3' | 5'-GTTCTCCGATCACATTCAGCAA-3' |
| RCOR1 | 5'-AACTGGCAAGACGCAGTCAA-3' | 5'-GTTGGGCAAATCAGCCAATGA-3' |
| CD36 | 5'-GGCTGTGACCGGAACTGTG-3' | 5'-AGGTCTCCAACTGGCATTAGAA-3' |
| IQSEC2 | 5'-GAGAAGGAAGCGGGCTATTCG-3' | 5'-TCCACACTGACTGTTCTGGAA-3' |
| ECHDC3 | 5'-CTGGACGGCATAAGGAACATC-3' | 5'-GACTTTCAGATCGTTGCTGTCA-3' |
| PSG3 | 5'-TGGGCCTGCATACAGTGGAC-3' | 5'-CTGGAGATGGAGGGCTTGGG-3' |
| AMMECR1 | 5'-TCTGCTCAGTGTCTCTGCTCA-3' | 5'-CGGTGCGTTTTGATCCTTTTTCA-3' |
| POMT1 | 5'-ATGGCCTACCAGATAGTGTTGG-3' | 5'-TGAGTGATGAGAGCATTCTCGAT-3' |
| ILKAP | 5'-CTAGCAGTGGCGATTCAGGTT-3' | 5'-TCACCGAAGAGGCTTTACAAAC-3' |
| PTCD1 | 5'-GAGCGTCAGGAAAACACGG-3' | 5'-AGGAGTATTTGTCAGAGAGGGTC- |
| ACTL8 | 5'-GAACATCGTGAACTACCTACCG-3' | 5'-CAAGGGTGTCTCCGTGATGAT-3' |
| ORC2L | 5'-GCTTCAGACAAGGTTCAACCG-3' | 5'-CTGTGCAACCCCTTCATCATC-3' |
| RGS17 | 5'-CAGAGGCCCAACAACACCTG-3' | 5'-TGTGGGTCTTCCCGCATTTT-3' |

| | | |
|---|---|---|
| SIAH1 | 5'-AGCCGTCAGACTGCTACAG-3' | 5'-AAAAGACTCGCCAAGTCATTGT-3' |
| ADCY9 | 5'-CAAGTTCGACTCGGTGAACCT-3' | 5'-CCGATGTAGAAGAGCGCATAC-3' |
| FOXN3 | 5'-ACTCTGACATGCCCTACGATG-3' | 5'-TCTGACTCCTCTCTTTGTCCAC-3' |
| MAP4K2 | 5'-GACACGGTCACGTCCGAAC-3' | 5'-ATTCCTGAGGTAGCTGCCAAT-3' |
| ZFX | 5'-TTGCTGAAATCGCTGACGAAG-3' | 5'-GCAATCGGCATGAAGGTTTTGAT-3' |
| PLEKHA9 | 5'-TGGTAAAACATTGCGGCAACA-3' | 5'-CCCTCTGCATCCCAATACTGAAA-3' |
| MRPS7 | 5'-GAGGAGAAATATGTTCGGGAGC-3' | 5'-CGTTCGATGGTTGCCTGTT-3' |
| EGFL7 | 5'-TGAATGCAGTGCTAGGAGGG-3' | 5'-GCACACAGAGTGTACCGTCT-3' |
| MYBPH | 5'-AGTATCAGAGTCCACCAGAGAAG- | 5'-TGCAGGCTTAGTGGCTGTG-3' |
| RHCE | 5'-CATCTCCGTCATGCACTCCAT-3' | 5'-TGAGTTCCCCAATGCTGAGGA-3' |
| PPP2R1A | 5'-TAGACGAACTCCGCAATGAGG-3' | 5'-CACCAGGGTAGTGAAGGTTCC-3' |
| APOH | 5'-CCCAAGCCAGATGATTTACCAT-3' | 5'-ACAGTCCTGTGAGAGGGCA-3' |
| USP10 | 5'-ATTGAGTTTGGTGTCGATGAAGT- | 5'-GGAGCCATAGCTTGCTTCTTTAG-3' |
| CASP6 | 5'-ATGGCGAAGGCAATCACATTT-3' | 5'-GTGCTGGTTTCCCCGACAT-3' |
| WISP3 | 5'-GGGCACTGGACCATTAGATACA-3' | 5'-TGAGTAGTCACAATACAGCCCT-3' |
| MKNK1 | 5'-AGATGGGCAGTAGCGAACC-3' | 5'-AGCAATTCAGAGGTCAGCTTG-3' |
| SNX11 | 5'-TTTTGGTGTAGGATGTCGGAGA-3' | 5'-GGCTTTGCTGTTGGTATGGAG-3' |
| RALBP1 | 5'-ATGGAAGGCTTAACGCTGAGC-3' | 5'-CTGTAGGACCACGTCGTTCG-3' |
| EBF2 | 5'- | 5'-CGACATTAGCGTCCACCACTC-3' |
| MAP2K7 | 5'-CCACGTCATTGCCGTTAAGC-3' | 5'-GCACGATGTAGGGGCAGTC-3' |
| OSBPL9 | 5'-CCAGAACCTGTTCAGTTGTGT-3' | 5'-ATGATGTCCCAAGGTAGCCAA-3' |
| TAAR5 | 5'-ACCAGGTGAATGGGTCTTGC-3' | 5'-TCCCTAGCACGATAATCAGCA-3' |
| LRRFIP2 | 5'-CCGATGGCACTCCCAATGG-3' | 5'-GCCTTACATCTAATGGCCCTTC-3' |
| MAN2A2 | 5'-TGTGTGGCAGTCTTCTCGC-3' | 5'-CGGAGTCCTTGATATGGCTGAT-3' |
| TPRKB | 5'-CTGGACCTATTTCCCGAATGC-3' | 5'-TTGTTTGCTGCCACAAGTATCT-3' |
| VPS26A | 5'-TTCAGGAAAGGTAAACCTAGCCT- | 5'-ATTGGCACCGATGTAAGATTCAT-3' |
| CPEB1 | 5'-GATGCAAATGACTTGTGCCTTG-3' | 5'-GGCTGAGGAATCTGAGTCCTG-3' |
| DDIT3 | 5'-GGAAACAGAGTGGTCATTCCC-3' | 5'-CTGCTTGAGCCGTTCATTCTC-3' |
| ADA | 5'-GCCTTCGACAAGCCCAAAGTA-3' | 5'-CTCTGCTGTGTTAGCTGGGAG-3' |
| CAV1 | 5'-GCGACCCTAAACACCTCAAC-3' | 5'-ATGCCGTCAAAACTGTGTGTC-3' |
| TUBA3C | 5'-AGGAGTCCAGATCGGCAATG-3' | 5'-GTCCCCACCACCAATGGTTT-3' |
| CTNNBIP1 | 5'-CCTATGCAGGGGTGGTCAAC-3' | 5'-CGACCTGGAAAACGCCATCA-3' |
| C1RL | 5'-TACCCAGAGCCGTATGGCAA-3' | 5'-GAACCGACGAATGAGATTGTGA-3' |
| SP1 | 5'-TGGCAGCAGTACCAATGGC-3' | 5'-CCAGGTAGTCCTGTCAGAACTT-3' |
| SACM1L | 5'-GCTGAAGCTGCATATCACACC-3' | 5'-ACGGTCAATGGTAAGTACGTCA-3' |
| C20orf59 | 5'-CACCTCGGGGATCGGATTG-3' | 5'-GCAGGGAAGTAAACCCCTTGG-3' |
| CACNA1I | 5'-GGAGCTGATCCTCATGTCCC-3' | 5'-CACGGGTTGCACACCATCT-3' |
| PLEKHB1 | 5'-CTGGAAGCGGAACTGGTTTG-3' | 5'-GGCCGATCTTTATGTCACGGA-3' |
| ZNF12 | 5'-ACAGGTGTGAAACTCTACAAGTG- | 5'-CCTCAGGTGGGTCGTGAGA-3' |
| CNOT2 | 5'-ACAGCATGTTTGGTGCTTCAA-3' | 5'-CTTTGTTGCCCGTATAAACTTGC-3' |
| MXRA8 | 5'-GCAACCTGCACCATCACTACT-3' | 5'-CCACCTGTTGAGCCTCCTC-3' |
| PZP | 5'-GGAGAAGGACTTATTCCACTGTG- | 5'-ATCTTGCGTAGGCCCCTTTAT-3' |
| IKZF3 | 5'-AAAGTGCAGCGGTTTTGAATG-3' | 5'-AGCTGTAAGGGATTTCAGGCT-3' |
| MIP | 5'-CTGTGAGGGAGCATTTGGTG-3' | 5'-TGTGCAACATTAGAGCTTGAGTC-3' |

| | | |
|---|---|---|
| FAM128B | 5'-TCCCGATGGCTCTCTCTGAAA-3' | 5'-TCTCCACGCCTTCTCAGGAT-3' |
| ZNF80 | 5'-TCCCAGGGACCAAGTAAAGAC-3' | 5'-CCTCATGGGTCGAACGAAGTC-3' |
| ASB1 | 5'-GGCTCCGCAGGTCGTAATC-3' | 5'-CCCGACGTAAGCTGCATCAT-3' |
| OSBP2 | 5'-TCGTCGCTGTTCACGGTTG-3' | 5'-CGGACACAGGTTCCGATCTTG-3' |
| OLIG2 (H) | 5'-CCAGAGCCCGATGACCTTTTT-3' | 5'-CACTGCCTCCTAGCTTGTCC-3' |
| OLIG2 (M) | 5'-GGCGGTGGCTTCAAGTCAT-3' | 5'-CATGGCGATGTTGAGGTCG-3' |
| SOX2 (H) | 5'-GCCGAGTGGAAACTTTTGTCG-3' | 5'-GGCAGCGTGTACTTATCCTTC-3' |
| SOX2 (M) | 5'-GCGGAGTGGAAACTTTTGTCC-3' | 5'-GGGAAGCGTGTACTTATCCTTCT-3' |
| POU3F2 (H) | 5'-CGGCGGATCAAACTGGGATTT-3' | 5'-TTGCGCTGCGATCTTGTCTAT-3' |
| POU3F2 (M) | 5'-GCAGCGTCTAACCACTACAGC-3' | 5'-GCGGTGATCCACTGGTGAG-3' |
| MYC(H) | 5'-ACTCTGAGGAGGAACAAGAA-3' | 5'-TGGAGACGTGGCACCTCTT-3' |
| MYC(M) | 5'-ATGCCCCTCAACGTGAACTTC-3' | 5'-GTCGCAGATGAAATAGGGCTG-3' |
| IL11 (H) | 5'-CGAGCGGACCTACTGTCCTA-3' | 5'-GCCCAGTCAAGTGTCAGGTG-3' |
| IL11 (M) | 5'-GCGCTGTTCTCCTAACCCG-3' | 5'-GCGCTGTTCTCCTAACCCG-3' |
| GAPDH | 5'-ACCCAGAAGACTGTGGATGG -3' | 5'-TTCTAGACGGCAGGTCAGGT-3' |
| ACTIN | 5'-TGAAGTGTGACGTGGACATC-3' | 5'-GGAGGAAGCAATGATCT-3' |
| 18S | 5'-TACCACATCCAAGGAAGGCAGCA- | 5'-TGGAATTACCGCGGCTGCTGGCA-3' |
